# Topological properties accurately predict cell division events and organization of *Arabidopsis thaliana’s* shoot apical meristem

**DOI:** 10.1101/2021.10.05.463218

**Authors:** Timon W. Matz, Yang Wang, Ritika Kulshreshtha, Arun Sampathkumar, Zoran Nikoloski

## Abstract

Cell division and the resulting changes to the cell organization affect the shape and functionality of all tissues. Thus, understanding the determinants of the tissue-wide changes imposed by cell division is a key question in developmental biology. Here, we use a network representation of live cell imaging data from shoot apical meristems (SAMs) in *Arabidopsis thaliana* to predict cell division events and their consequences at a tissue level. We show that a classifier based on the SAM network properties is predictive of cell division events, with validation accuracy of 82%, on par with that based on cell size alone. Further, we demonstrate that the combination of topological and biological properties, including: cell size, perimeter, distance, and shared cell wall between cells, can further boost the prediction accuracy of resulting changes in topology triggered by cell division. Using our classifiers, we demonstrate the importance of microtubule mediated cell-to-cell growth coordination in influencing tissue-level topology. Altogether, the results from our network-based analysis demonstrates a feedback mechanism between tissue topology and cell division in *A. thaliana*’s SAMs.

**Summary statement:** we use a network representation of live cell imaging data from SAMs in Arabidopsis thaliana to predict cell division events and their consequences at a tissue level.

## Introduction

The adjacency of cells, specifying the tissue topology, defines the organization of cells and affects function of organs in multicellular organisms. Therefore, deciphering the organizational principles of cellular connectivity networks are fundamental to improve our understanding of the development of multicellular organisms. The shoot apical meristem (SAM) of plants is a highly organized structure composed of continuously proliferating cells that differentiate and give rise to all aerial organs and is under the control of an intricate signaling network influencing plant growth and response to different stimuli. The SAM epidermis in plants serves as an excellent system to identify organizational principles of cellular connectivity networks (Varner and Lin, 1989).

Since the cells in the SAM are glued to each other by a rigid cell wall, changes in the topology of SAMs are only brought about by cell division events. Cell division in plants is a cell-size-dependent, cell autonomous process (Jones et al., 2017), and crossing multiple checkpoints allows the final transition towards cell division (Veylder et al., 2007; Qi and Zhang, 2019). Willis et al. (2016) recently showed that initial cell size at birth influences the increase in size (sizer model), even though there seems to also be a component of constant size increase (adder model) in the shoot apical meristem (SAM) of *Arabidopsis thaliana*. This study has hinted at the possibility that a combination of both models may best describe cell division (see D’Ario and Sablowski (2019) for a comparison of models). Although size-dependent cell division seems to be independent from position and cell to cell contact (Willis et al., 2016), recent study of Jackson et al. (2019) has pointed out that dividing cells display higher centralities in the network representation of the *A. thaliana*’s SAM; however, this observation was not sufficient to accurately predict cell division from network properties alone.

Since biochemical and physical signals are transmitted across tissues and affect cell division, growth, and morphology in a spatio-temporal fashion, the question arises of how tissue topology could influence such processes to help the plant respond to a variety of stimuli. In the context of physical signals, the ability of plant cells to respond to growth driven mechanical signals requires the activity of microtubule severing protein KATANIN (Uyttewaal et al., 2012). It has been shown that the lack of mechanical feedback, as in the *katanin1-2* mutant, results in changes to the topological features as a consequence of modified cell shape (Jackson et al., 2019). Therefore, this mutant can be employed to test if topological features are indeed relevant for cell division and related processes.

This question can be readily addressed due to the availability of plant lines expressing stable fluorescence reporters that allow for monitoring cellular outlines in combination with confocal imaging techniques (Reddy et al., 2004). In addition, the combination of user-friendly tools for accurate segmentation, like MorphoGraphX (Barbier de Reuille et al., 2015), with different machine learning (Bhavsar and Panchal, 2012; Pisner and Schnyer, 2020) and deep learning techniques (Camacho et al., 2018) has led to massive advances in the analysis of high-throughput imaging data. Further, the analysis of imaging phenotypes has been facilitated by adopting the network paradigm (Breuer et al., 2017; Nowak et al., 2021). To this end, topological features have been employed in cell wall placement models for dividing cells, by using the degree (i.e. number of neighbors) in combination with a spring based model (Gibson et al., 2011) or other individual topological features (Jackson et al., 2019). It has been shown that some of these individual topological features can better predict the placement of certain cell walls compared with more traditional approaches (Jackson et al., 2019), such as: dividing cells using the shortest wall placement, generalized Errera’s rule (Besson and Dumais, 2011) or by minimization of tensile stress in other models (Louveaux et al., 2016). Although these models present an important step to solve the problem of cell wall placement, each model underperforms on some cells in the central region of the SAM (Shapiro et al., 2015; Jackson et al., 2019).

Although there are attempts of combining network properties with imaging data from SAMs, minor progress has been achieved in predicting individual cells divisions in this plant tissue. Here, we provide a network-based perspective to model cell division and cell wall placement in the SAM of *Arabidopsis thaliana*, a well-established system for studying cell division. To this end, we combine network-based analysis of live cell imaging data with classifiers that allow us to simulate tissue-wise topological changes of the *A. thaliana* SAMs and test these classifiers independently on the *katanin1-2* mutant.

## Results

### Topology and surface area accurately predict cell division events

The question of whether division of a cell embedded in a tissue is driven by the topology of the neighboring cells, the area of the cell, or combination of the two is still open. To address this question, we imaged SAMs of five *A. thaliana* expressing a plasma membrane reporter (pUBQ10:acyl-YFP) every 24 hours over five days using confocal microscopy (Figure 1A). First, we manually determined the number of dividing and non-dividing cells between two consecutive time points in the central zone of the SAM. We defined the central zone of a SAM as the area covered by a circle of 30 μm radius around the highest point in the analyzed SAM (Figure 1B), and found that 24.3%±3.5% of cells divided per tissue between two successive time points, with a total number of 329 dividing cells and 896 non-dividing cells (Figure 1B).

**Figure 1.**
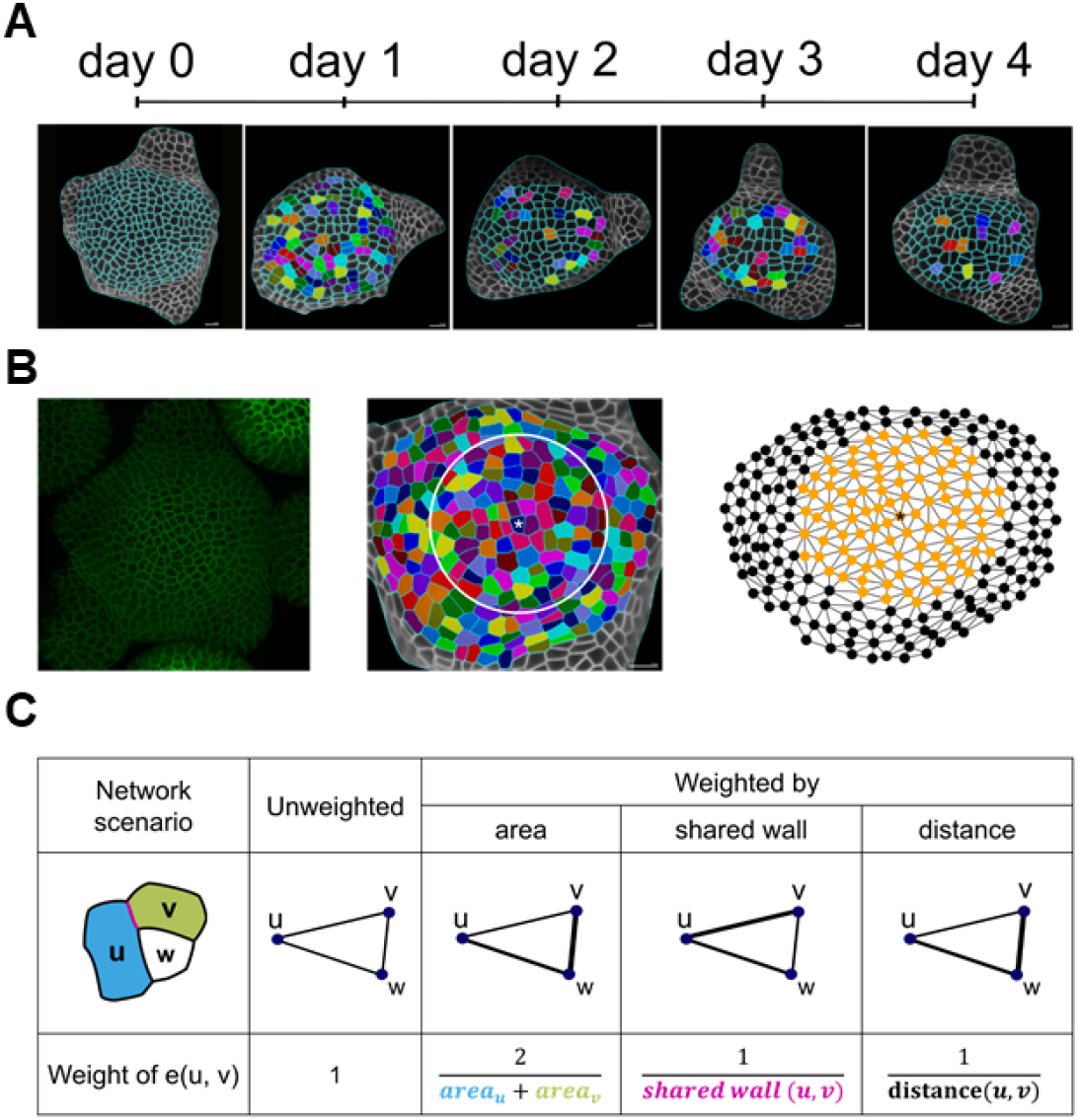
Feature generation from three-dimensional (3D) images of the shoot apical meristem (SAM). (A) The surface of A. thaliana SAM is imaged every 24 hours over four days. Pairs of dividing cells, depicted with the same colors, are determined manually (see Methods). (B) The 3D images of SAMs are converted to 2½D surfaces by employing MGX (de Reuille et al., 2015) (left panel). The surface is abstracted by its topology, capturing the connectivity of neighboring cells in a radius of 30 μm (white circle) around the central cell, marked with * (center panel). The topology of the analyzed cells inside the circle is colored in orange (right panel). Two nodes are connected by an edge if the cells they represent share cell wall. (C) Four different network scenarios are considered: (i) unweighted edges and edges weighted by (ii) area, (iii) shared cell wall, and (iv) distance, illustrated for the case of three cells u (blue), v (green), and w (white). In the unweighted network scenario, all edge weights have a value of one. The edge weight for the network weighted on area, shared wall, and distance is the inverse of the mean cell areas of u and v, of the shared cell wall area (magenta), and of the inverse distance of the center of mass for the graph weighted on the distance (black). The weights of the edge e(u, v) in the four scenarios are illustrated with different line widths.

Next, we represented the topology of the central zone as a network, in which every node corresponds to a cell and two nodes are connected by an edge if the cells share cell wall. For each cell we calculated 16 properties, referred to as topological features (Supplementary Table 1), in an unweighted network, in which every edge is of weight 1. We also applied different edge weights based on the mean surface area, shared cell wall, and distance of the cell centroid between two nodes representing those cells (Figure 1C). In addition, we considered the surface area of each cell in the central zone as a biological feature.

Previous studies have shown that there exists a critical cell size threshold for cell division in the SAM of *A. thaliana* (Jones et al., 2017). To show that topological features capture information distinct from that provided by the cell surface area, we calculated its Pearson correlation with the topological features (Supplementary Figure 2, Supplementary Table 2). Using the network with edges weighted based on the cell surface area, we found that betweenness centrality, a measure for the relative number of shortest paths passing through a node, exhibited the highest correlation of 0.71 to the surface area. Nevertheless, the absolute value of the correlation with surface area was smaller than 0.5 for 63% of the features (97% of features showing correlation smaller than 0.7). Therefore, topological features in the considered network scenarios carry information that is different from that obtained by the cell surface area alone. To further show the predictive power of the classifiers trained on the topological features, we considered two reduced feature sets that only included features with absolute value of the Pearson correlation coefficient (r) smaller than 0.5 and 0.7, respectively (Supplementary Figure 2). In such a way, we aimed to remove bias due to consideration of features which may, to a certain extent, include information about surface area.

As a result of these considerations, we trained six classifiers based on non-linear support vector machines (SVMs) with Gaussian kernel ((Bhavsar and Panchal, 2012)) to predict cell division based on: all topological features (topo), surface area alone, the combination of topological features and surface area (topo + area), topological features with low absolute value of correlation with surface area (r < 0.7 and r < 0.5), and on unweighted topological features. To this end, we selected an equal number of dividing and non-dividing cells from four SAM for training the SVMs, to ensure balancing of cell labels. We kept the data from the remaining, fifth SAM, as a testing set (Methods). Further, we partitioned the 502 selected cells into training and validation sets composed of equal numbers of dividing and non-dividing cells, and used five-fold cross validation to train the classifiers (see Methods).

While the training accuracy of the SVM using only the surface area was 79.4%, the training accuracy solely based on topological features was significantly higher, at 88.7% (11.0% higher; p-value=0.0011, one-way ANOVA); this was also the case when the combined set of topological features and surface area was used, with training accuracy of 86.3% (8.3% higher; p-value=0.0101, one-way ANOVA). However, we observed no difference in the validation accuracies for the three types of SVMs (∼81%). For the test SAM, the classifier based on the combination of topological features and surface area exhibited the best performance, with an accuracy of 78.9%, followed by the SVM that considered the topological features (76.9%) and the surface area (72.4%) alone (see Figure 2, Supplementary Table 3). The area under the curve (AUC) of the receiver operating characteristic (ROC) - curve, used as another measure of performance, showed similar trends (Supplementary Figure 3A, Supplementary Table 3).

**Figure 2.**
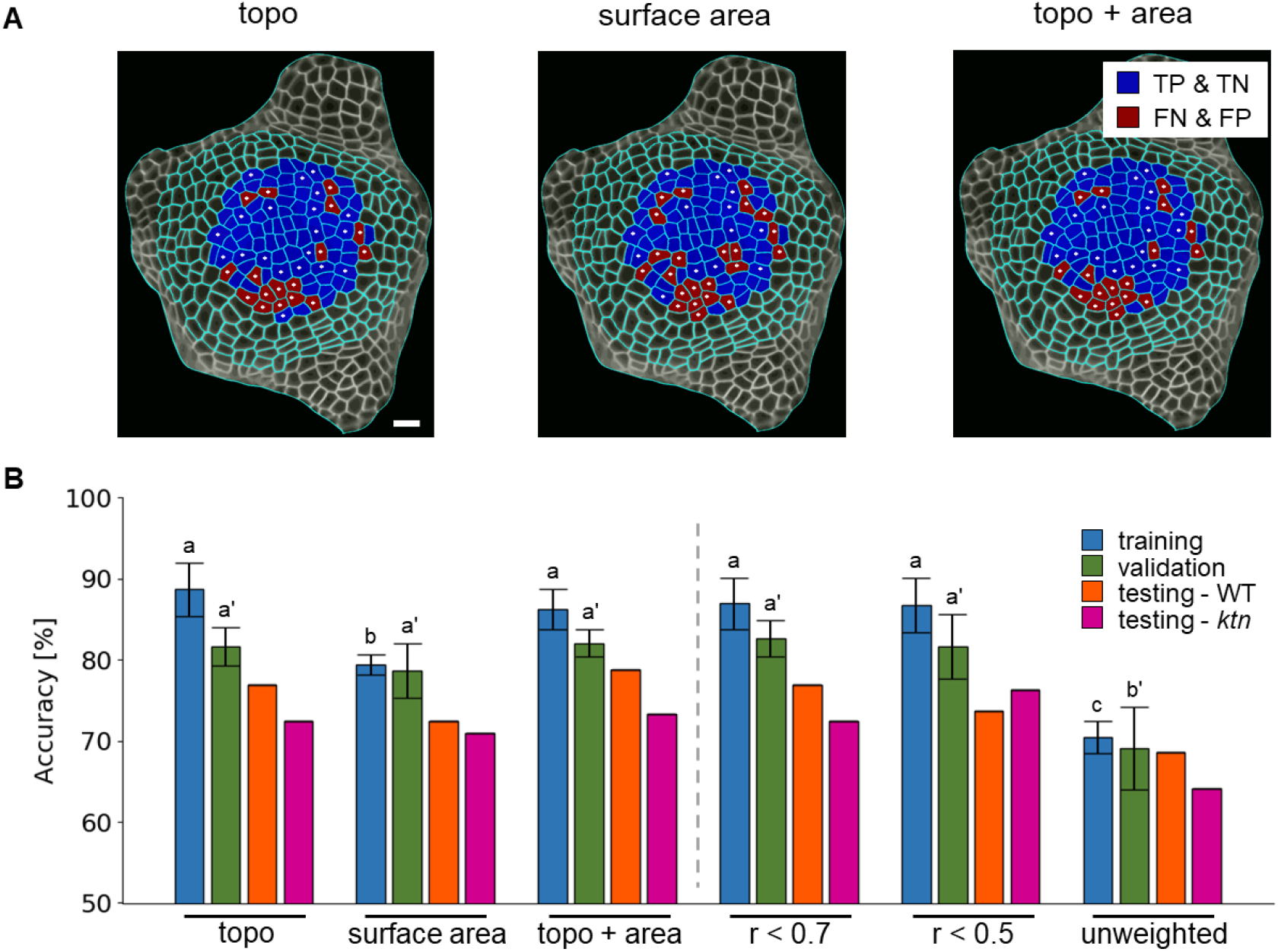
Surface area and topology-based features generate different predictions for cell division in the SAM. (A) Comparison of predicted and observed labels on one test plant tissue from Figure 1A day 0 highlighting the difference between predicted and observed division events. The predictions are from classifiers trained on different feature sets: combined topological features (topo), surface area, combined topological features and surface area (topo + area; from left to right) with the coloring scheme of correct predictions in blue and wrong prediction in red. The combined topological features include 16 centrality measures (see Methods) calculated based on the four network scenarios (see Figure 1C). Dividing cells are marked with a white star. Scale bar is 10 μm. (B) Accuracy of the support vector machine classifier to predict cell division for the training (blue), validation (green), and testing of wild type (orange) and *ktn* mutant (purple) data sets using: topo, surface area, and topo + area, reduced set of topological features that show an absolute Pearson correlation coefficient with surface area smaller than 0.7 or 0.5 (r < 0.7 and r < 0.5), as well as only the topological features derived from the unweighted network scenario (unweighted). The performance on the training and validation set is determined from five-fold cross-validation with mean and the standard deviation shown as error bars. Different letters indicate significance between groups using one-way ANOVA with Tukey’s pairwise comparison: p-value < 0.05. Statistical testing for differences of classifier performance for the training and validation sets was conducted separately (letter without and with apostrophe, respectively). N_WT_ = 5 plants, 4 time steps (4 plants for training-validation and 1 plant for testing); N_*ktn*_ = 3 plants, 3 time steps; n_WT_ = 502 and 156, train-validation and test cells respectively; n_*ktn*_ = 334 (balanced data).

The removal of topological features that were highly correlated with area does not significantly change the validation accuracy (Figure 2, Supplementary Table 3). Moreover, using only the topological features from the unweighted network scenario (Figure 1C), resulted in 16.7% smaller accuracy of the classifier on the validation set in comparison to that based on all topological features (p-value < 0.001, one-way ANOVA). However, the classifier based on the features from the unweighted topology only performs slightly worse (relative difference in accuracy of 5.5%) compared with the classifier based on the surface area (Figure 2, Supplementary Table 3). Inspection of the learning curves showed that the classifiers did not suffer from high bias and variance and that the training set was sufficiently large (Supplementary Figure 4). Therefore, we concluded that topological features led to marginally improved performance in predicting cell division compared to the surface area alone, while the performance could be further increased by the combination of both feature types.

To further corroborate the biological relevance of these findings, we randomly permuted the labels and retrained the classifiers, repeating this procedure 1000 times for each feature set (see Methods). All but the classifier based on surface area were able to partially predict training data. However, no classifier was able to generalize on the validation or test set, exhibiting accuracies expected by chance (Supplementary Figure 5A). Therefore, the classifiers trained on the randomized labels demonstrated that the used features capture information important for classification of dividing and non-dividing cells in the 1-day long intervals.

Independent testing of the trained SVMs with data from the *katanin1-2* (*ktn*) mutant to predict cell division events showed reduced accuracy in comparison with data from the wild type (WT) for all classifiers. The relative difference between the test accuracies for WT and *ktn* was smaller for the classifier trained on surface area (2.0%) compared with all topological (6.0%) as well as the topological and biological features combined (7.2%). Removing the features with Pearson correlation to surface area higher than 0.7 led to relative decrease of 6.0% between the test accuracies for WT and *ktn* (Figure 2). While these findings demonstrate the importance of surface area as a determinant of cell division, they also support the claim that topology plays an important role in predicting cell division events.

### Combination of topological and biological features enables recreation of the local topology after cell division

To examine whether properties derived from the tissue connectivity network as well as biological properties (i.e. cell size and perimeter, distance, and shared cell wall between cells) are predictive in the time-dependent connectivity of daughter cells, we trained classifiers based on SVMs (with Gaussian kernel) to predict which of the cells adjacent to a dividing (parent) cell are neighbors of the divided (daughter) cells. We distinguished neighbors that were only adjacent to one daughter cell: adjacent to the daughter cell closer to the SAM center are labelled as class 0, while those adjacent to the daughter farther from the center were classified as class 1; neighboring cells adjacent to both daughter cells were considered to be of class 2 (Supplementary Figure 1C).

To predict changes of cell divisions based on local topology from the data collected at 1-day time interval, we first determined all neighbor-parent pairs and then predicted the adjacency of the neighbor to the daughter cells. To this end, we considered the topological features as well as biological properties of the parent and the neighbor cells in the preceding time point. We used the topological features of the neighbor cell to distinguish neighbor-parent pairs in which a neighbor is adjacent to two dividing cells. We also included the difference between the topological features of the neighbor and the parent, thus generating 100 unique topological-based properties for each neighbor-parent-pair (Supplementary Figure 1B). For the biological feature set, we extracted the surface area and perimeter of both the parent cell and neighbors as well as their shared cell wall and distance between the centers of mass. For the combined features, we concatenated both topological and biological features. Given a parent cell, we determined the class of its neighbors in the next time point by aligning the tissues between 1-day time intervals manually, and determining their adjacency of the neighbor with respect to the daughter cells (see Methods and Supplementary Figure 1B).

We excluded neighbor-parent-pairs for which the neighbor also divided in the considered 1-day interval to avoid bias due to guessing which cell divides first (Figure 3A). Following this procedure, we created 1638 neighbor-parent pairs with 546 representatives (balanced classes) in each of the three classes (0, 1, and 2) from five different SAMs, tracked every 24 hours over 5 days. The data was split into three parts: training, validation, and test data, such that the SAM of one plant was kept as test data, while the rest of the plants were used in a nested five-fold cross validation for training the SVM.

**Figure 3.**
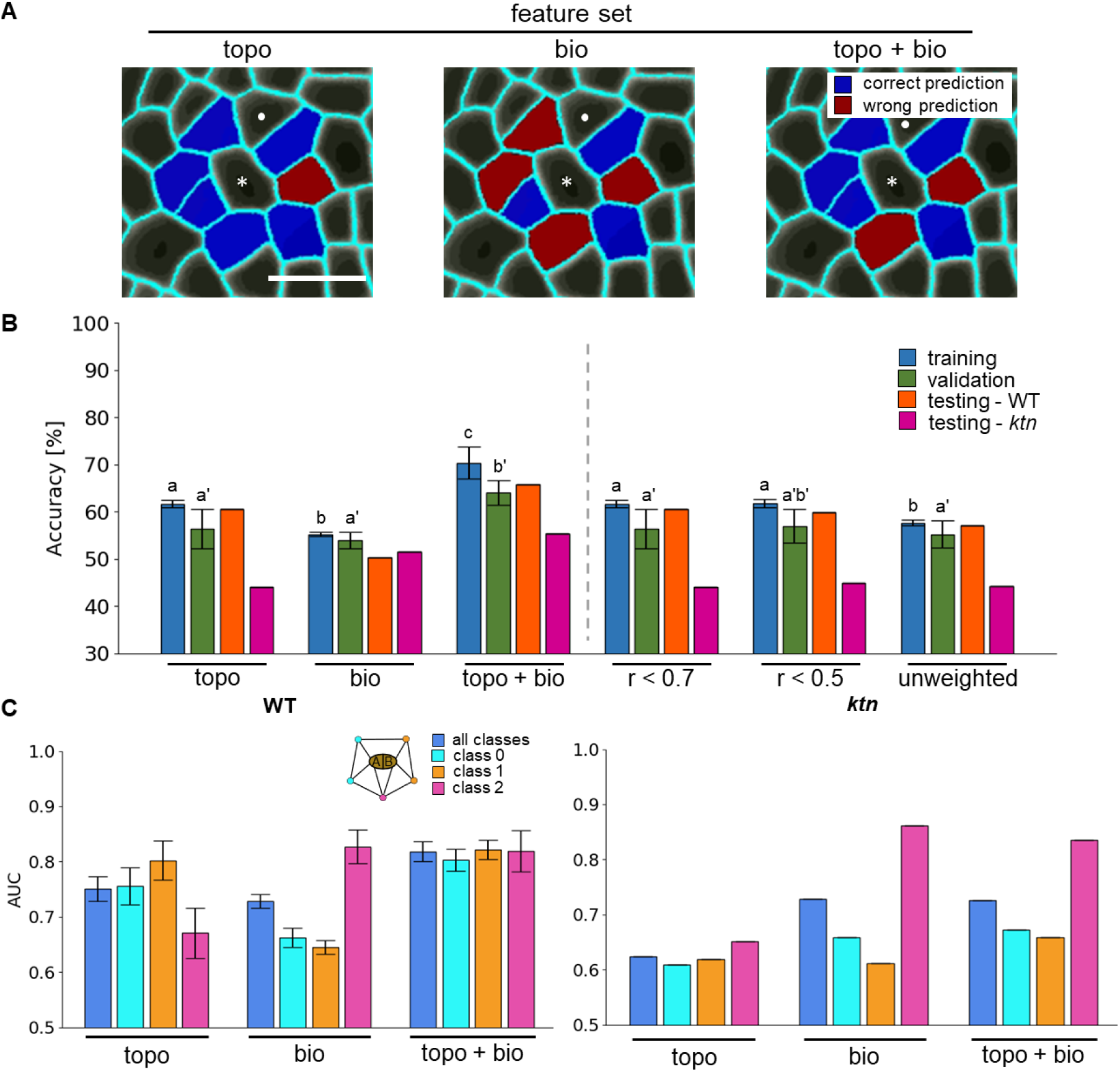
Topological and biological features are required for accurate prediction of the local neighborhood after cell division in the SAM. (A) Comparison of predicted and observed local neighborhoods on one local topology from Figure 1A, day 0. The predictions are made with classifiers trained on different feature sets: combined topological (topo, including features of the four network scenarios, see Figure 1C, Supplementary Figure 1), biological (bio, including area, perimeter, shared cell wall, and distance), and combined topological and biological features (topo + bio; from left to right) with the coloring scheme of correct predictions in blue and wrong prediction in red. The combined topological features include 16 centrality measures (see Methods) calculated based on the four network scenarios (see Figure 1C). The central and neighboring dividing cells are marked with a white star and circle, respectively. No color is displayed on the dividing neighbor cell as all dividing neighbors were removed. Scale bar is 10 μm. (B) Accuracy of classification on the training (blue), validation (green), and testing of wild type (orange) and *ktn* mutant (purple) and (C) Area under the curve (AUC) of the ROC based on the wild-type validation (left) and *ktn* test data (right) for the classification of all classes (blue), for the class of neighbors adjacent to the daughter cell (cell A, see legend in B) closer to the SAM center (denoted as class 0; cyan), adjacent to the daughter (cell B, see legend in C) farther from the center (denoted as class 1; orange), or adjacent to both cells (denoted as class 2; magenta). The classifiers are based on topo, bio, topo + bio, reduced set of topological features that show an absolute Pearson correlation coefficient with all biological features smaller than 0.7 or 0.5 (r < 0.7 and r < 0.5), as well as topological features derived from the unweighted network scenario (unweighted). The performance on the training and validation set is determined from five-fold cross-validation with mean and the standard deviation shown as error bars. Different letters indicate significance between groups using one-way ANOVA with Tukey’s pairwise comparison (p-value < 0.05). Statistical testing for differences of classifier performance for the training and validation sets was conducted separately (letter without and with apostrophe, respectively). N_WT_ = 5 plants, 4 time steps (4 plants for training-validation and 1 plant for testing); N_*ktn*_ = 3 plants, 3 time steps; n_WT_ = 1317 and 312, train-validation and test cells respectively; n_*ktn*_ = 888 (balanced data).

The training and validation accuracy was best for the SVM based on the topological features combined with biological features, at 70.4% and 64.0%, respectively. The topological and biological features alone showed 12.7% or 17.0% reduction in validation accuracy compared to the combined classifier, and similar reduction in training accuracies. Regarding the performance on the test set, the combined classifier performed best, with an accuracy of 65.7%, followed by the classifier based on topological features alone with 60.6%, and that using biological features alone, with the worst accuracy (equivalent to guessing) of 50.3% (Figure 3B, Supplementary Table 4).

Investigating the area under the ROC-curve (AUC) measure for individual classes highlighted the differences between the two classifiers trained on topological or biological features alone: The SVM based on the topological features showed better performance for the neighbors only adjacent to one cell (class 0 and 1) in comparison with the classifier based on the biological features (i.e. relative increase of 10.7% and 23.9% for class 0 and 1, respectively). In contrast, the SVM based on the biological features performed 22.6% better for neighbors adjacent to both daughter cells in comparison with the classifier based on the topological features. Combining both feature sets improved the average AUC on the validation data of the classifier by 9.4% and 13.0% (relative increase compared to topological and biological features alone, respectively) while retaining high performance for all classes (Figure 3C, Supplementary Figure 6). Investigating the reduced topological feature set (i.e. removing features with Pearson correlation coefficients larger than 0.5 or 0.7 with any biological feature) as well as only considering unweighted features resulted in almost identical training, validation, and test accuracies compared with all topological feature trained classifiers. The training accuracy of the unweighted set showed a slight relative reduction of 7.0% (p-value: 0.013, one-way ANOVA) (Figure 3A, Supplementary Table 4). These findings indicated the importance of both topological as well as biological properties to predict local topology after a division event.

To further corroborate the biological relevance of these results, we randomly permuted the labels and retrained the classifiers, repeating this procedure 1000 times for each feature set. While the resulting classifiers showed performance better than expected at random on the training data with the three sets of features, they did not generalize well and exhibited accuracy on the validation set similar to expected by chance (Supplementary Figure 5B). Further, we investigated the more difficult scenario of including neighbor-parent-pairs whose neighbors also divide and repeat the topology prediction procedure. Here, we found similar performance to that on the training, validation, and test sets for all combinations of feature sets (Supplementary Table 4). Therefore, our findings demonstrated that the used features capture information important to classify changes in local topology predictions surrounding dividing cells in 24-hour intervals.

We tested how well the trained classifiers based on the wild-type data performed on the *ktn* mutant. With the data from the *ktn* mutant, we found a reduction in accuracy for all classifiers for local topology prediction, trained on the wild type data, except for those using the biological feature set (50.3% WT testing vs 51.6% *ktn* testing). All classifiers trained on topology-related feature sets showed strong relative reduction in test accuracy between WT and *ktn* (topo: 31.6%; topo + bio: 17.0%; r < 0.7: 33.3%; r < 0.5: 28.9%; unweighted: 25.5%; orange vs purple bars, Figure 3B). The reduced performance on *ktn* data of classifiers trained on topological features (feature sets: topo, topoAndBio) can be mainly attributed to the worse prediction of class 0 and 1 (cells being predicted to be adjacency to only one of the divided daughter cells, Supplementary Figure 1C). In contrast, the classifiers trained on biological features performed similarly on *ktn* test as those based on the WT test data (Figure 3B). These results highlight the importance of both topological and biological information in local topology rearrangement after cell division.

### Combined application of division event and local topology prediction enables to predict tissue topologies

To apply the classifiers and compare the resulting topologies, we used the data from the test plant and successively predicted division events and changes in local topology using classifiers trained on the combined biological and topological features (see following procedure in Figure 4A). We compared the predicted and observed topologies by investigating the unweighted topological features (Figure 1C, Supplementary Table 1) of non-dividing cells in the next time points of both scenarios. We selected non-dividing cells of both scenarios, i.e. predicted and observed, to pairwisely compare their unweighted topological features. We did not consider other network scenarios (see Figure 1C) since we would need to estimate the weights for the topology, adding a layer of uncertainty. Here, 126 of the possible 155 non-dividing cells in the observed topology were also non-dividing in the predicted topology. For these cells, we calculated the Pearson correlation coefficient (r) of all unweighted features between observed and predicted topologies, with the harmonic centrality showing the largest value of r = 0.80 and eight of 16 features having Pearson correlation larger than 0.5 (Figure 4D, blue bars). We compared the predicted and observed values of harmonic centrality of the non-dividing cells of the next time step, and found strong correlation (Figure 4E, Supplementary Figure 8).

**Figure 4.**
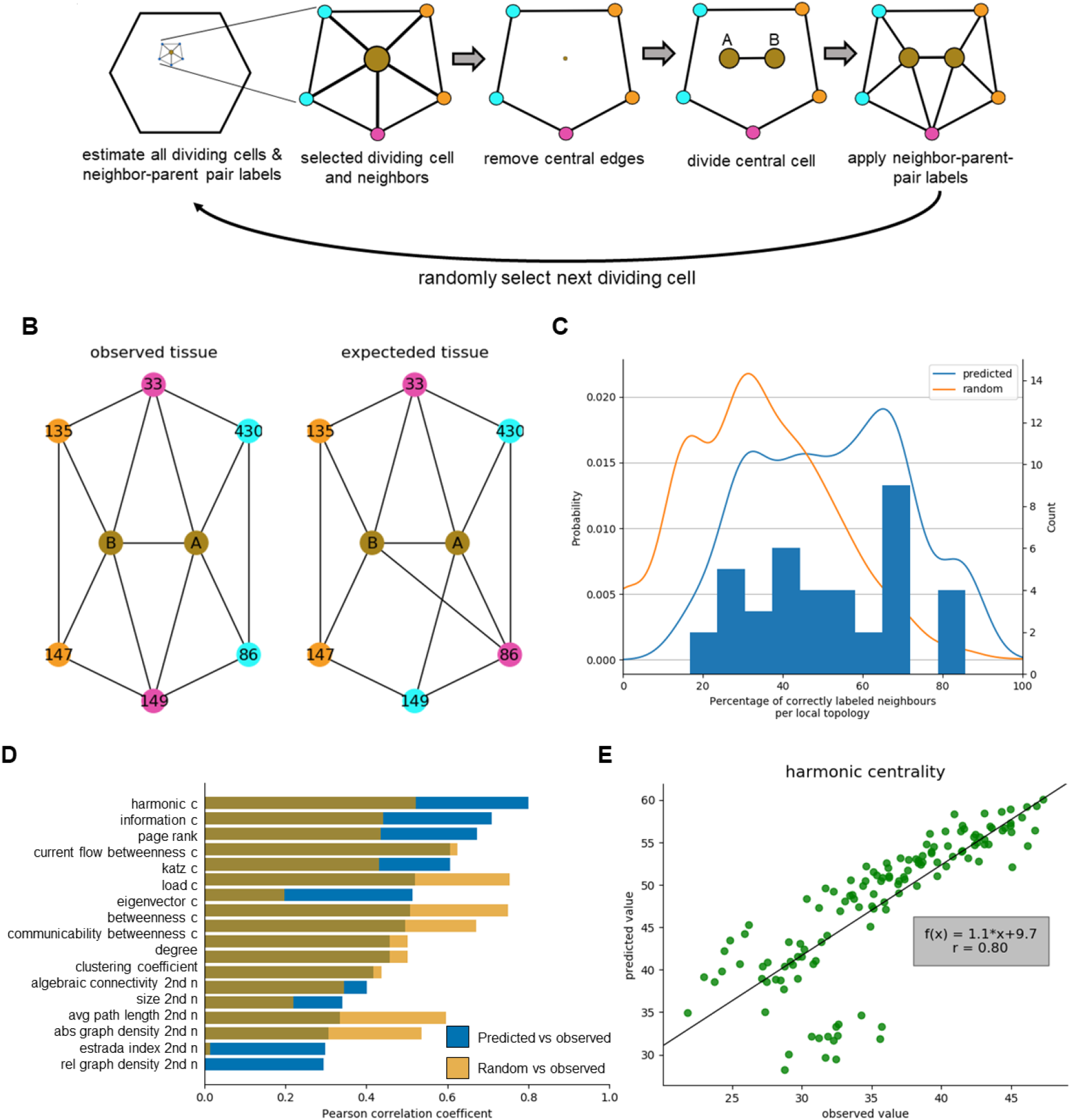
Concordance between observed and predicted topologies. The cell connectivity network of the test plant was predicted by applying classifiers for division event and topology prediction. (A) Illustration of the procedure applying the division and local topology classifiers to generate the topology of the next time point (24-hour time interval). After division and topology prediction, a dividing cell (dot in brown) is selected along with its neighbors (blue circles) and its adjacency relationship (edges, black lines) (left). The selected cell (predicted to divide) along with the edges incident to the corresponding node are removed and replaced by the divided daughter cells (A, B: representing the cell closer and farther away from the SAM center) that are adjacent to each other (three in the middle). The daughter cells are connected with their neighbors based on the prediction from the local topology classifier (right). The next dividing cell is randomly selected and the previous steps are repeated until all dividing cells are selected. (B) One example of the predicted local topology with an overall accuracy of 66.6% for the full local topology is compared with the observed local topology. The divided daughter cells “A” and “B” (brown) are adjacent with the predicted or observed cells (numbers display the same cells) coloring their respective parent-neighbor-class (cyan/orange: cell connected only with daughter A or B, respectively; magenta: cell adjacent with both daughter cells) (C) Histogram and density plot of the percentage of correctly estimated neighbors per local topology of cells predicted as dividing in the test plant (blue) are compared with the density plot of randomly assigning parent-neighbor-classes (orange). Difference between distributions is tested using Kolmogorov-Smirnov-Test, p-value < 0.01. N = 1 plant, 4 time steps, n = 39 local topologies. (D) The concordance between the observed and predicted topologies was quantified (blue) for non-dividing cells in both topologies by calculating and ranking the Pearson correlation coefficient based on 16 topological features from the unweighted networks (see Figure 1C; Supplementary Table 1). The procedure was repeated dividing all cells predicted to be non-dividing, randomly assigning classes to the neighbors, and calculating the correlation as described before (yellow). (E) The observed harmonic centrality is plotted against the predicted harmonic centrality for all non-dividing cells and the best linear fit (solid line) with its function f(x) and the respective Pearson correlation coefficient r is overlaid. N_WT_ = 1 plant, 4 time steps, n_WT_ = 126 cells not dividing in both observed and predicted tissue.

For comparison, given the same test plants observed topologies, we selected all predicted non-dividing cells to divide and randomly connected the neighbors with the divided cells reorienting their local neighborhood; we then repeated the correlation analysis of the resulting topology with the observed one. This “random propagation” scenario allowed us to construct and investigate the most opposite example to our predictions (Figure 4D, orange bars). Comparing the predicted and random propagations correlations shows that only five out of 16 topological features showed higher correlation in the random propagation. The random propagation showed the lowest correlating features and the trained classifiers showed the highest correlations with a total of eight being higher (Figure 4D, blue vs orange bars).

To further investigate the performance of the local topology prediction on the test plant, we calculated the percentage of correctly predicted neighbors for each cell dividing in the predicted and observed tissue (example in Figure 4B). The distribution of correctly labelled neighbors per dividing cell was significantly shifted towards higher accuracy when comparing the predicted and random topology predictions (Figure 4C).

## Discussion

The biochemical pathway of cell division control has been extensively studied (Dewitte and Murray, 2003), but only recently external cues have also been considered to understand the effect of cell divisions in a tissue context (Hartig and Beck, 2006; Shimotohno et al., 2021). It has been known that the outer epidermal cell wall resists most forces (Beauzamy et al., 2015) and, thus, division in the SAM outer-layer needs to serve both meristematic functions. This raises the question if cell division and their subsequent local topology rearrangement are affected by the tissue topology and if tissue topology contains sufficient information for their accurate prediction.

Based on our extensive network-based modelling, we showed that both surface area, as an approximation of cell size, as well as the characteristics of topology allow for prediction of cell division events in the central epidermal region of *A. thaliana* SAM, in contrast to earlier reports (Jackson et al., 2019). The cyclin-dependent kinase (CDK) G1 is known to bind DNA and serves as a ruler after cell division, allowing for size dependent division in *C. reinhardtii* (Li et al., 2016), while KIP-related protein 4 has a similar function in the *A. thaliana* SAM niche (D’Ario et al., 2021). Modelling cell division in the SAM of *A. thaliana* also revealed the importance of CDKs in G1-S and G2-M phase transition (Jones et al., 2017). Further, the work of Willis et al. (2016) showed that cell division events in SAMs of *Arabidopsis* treated with naphthylphthalamic acid, an inhibitor of auxin transport that generates naked meristem, are influenced by both cell size increase and a cell size threshold affecting cell division. Both models explain the importance of surface area in predicting cell division events, but they do not explain the importance of topological features. Here, the mechanical feedback loop, envisioned by the cells ability to react to changes in turgor pressure with MT and cell wall rearrangement affecting cell divisions (for detail of the feedback loop, see Sampathkumar (2020)) may serve as an explanation linking topology with the summed turgor and supracellular mechanical stress. Alternatively, the predictive ability of topological properties may result from long distance communication by different phytohormones (Shimotohno et al., 2021), or due to cell-to-cell communication by plasmodesmata (Kitagawa and Jackson, 2017).

However, not only the cell division, but also the cell wall positioning affects the tissue organization; a prime example is the effect of division patterning in lateral root initiation (Wangenheim et al., 2016). Our study relies on the adjacency of cells in the tissue topology, in contrast to other cell wall models, such as: the generalized Errera’s rule (Besson and Dumais, 2011), the spring-based model (Gibson et al., 2011), and the mechanical stress related model (Louveaux et al., 2016), that predict the placement of the cell wall based on the individual cell geometry. Our classifier employs the biological feature set composed of six cellular features, having limited information about the dividing and neighbor cell geometry, and allows for reliable prediction of the changes in the local topology. These local changes in the topology mirror the effect of the cell wall placement on the tissue. In addition, we showed that topological features alone sufficed to accurately predict local topological changes. While single topological properties were already used to estimate cell wall placement (Jackson et al., 2019), the percentage of dividing epidermal cells in this study was only 12% (total n=7/57 dividing and non-dividing cells) per tissue every 22h. In contrast, our results rely on experiments in which cells divided more regularly, with an average of 24% of dividing cells per tissue every 24h (total n=329/896 dividing and non-dividing cells), allowing us to train robust classifiers. We showed that the combination of both feature sets boosted performance of local topology reorientation prediction (Figure 3), indicating that the inclusion of multiple viewpoints of information available to cells needed to be involved to solve the problem of cell wall placement in the SAM. This raises the question how information of the topology is biologically transferred to cells, either via mechanical stress, hormones, or cell to cell communication with plasmodesmata.

To demonstrate the generalizability of the classifiers, we showed that they can be used to make accurate predictions for *ktn* mutants that are defective in mechanical feedback regulation. Our results indicated similar performance for the classifiers with biological feature sets from WT and *ktn*. In contrast, the classifiers trained on topological features showed reductions in performance in *ktn* compared to WT. This difference in performance is not due to differences in topological features, since the normalized features showed similar distributions (Supplementary Figure 9). These results suggest a potential role of KATANIN in linking sub- and supracellular mechanical stress, known to affect leaf epidermal cells (Eng et al., 2021) and KATANINs role in positioning of the preprophase band, spindle, and phragmoplast (Komis et al., 2017). In addition, the cell geometry of the *ktn* mutant differs compared to the WT and might also influence the topology. Therefore, the combination of network-based modelling with machine learning provides a method to screen SAMs under different conditions and mutants. More specifically, reduction in test performance of either the classifiers trained on surface area or on topological features compared to the wild type may hint to effects only disturbing function related to the cell cycle or to a topological effect (in the case of classifier trained on area or topological features being lower, respectively).

When combining division prediction and the resulting changes to the tissue, previous studies mostly focus on single cell division or propagating tissues based on division likelihoods using the number of neighbors (Gibson et al., 2011) or just using area as a fixed threshold (Sahlin and Jönsson, 2010; Alim et al., 2012), while our classifier incorporates more diverse tissue-level information. Here, we combined our best classifiers to predict future tissue topology using the combined topological and biological features. Although the results of this propagation of classifiers is promising, the careful inspection of the finding, particularly with respect to planarity and topological properties of the reconstructed topologies point out that further research should consider simultaneous modelling of cell neighborhoods of higher order to improve the reconstruction.

Furthermore, as information is not only be passed along the epidermis (L1-layer), the assessment of cell division events and their changes on the topology could be expanded beyond the epidermis of the SAM as we know that the L2- and L3-layer play a vital role in supporting the meristematic function through the feedback of CLAVATA 1, 2, and 3 and WUSCHEL (Schoof et al., 2000). Transferring the classifiers to other plant species, such as maize (that has only two distinct layers forming the SAM), may provide insights into how meristematic function can be conserved with fewer cells. As other tissues and organs are also experiencing mechanical stresses, hormone gradients and other transport related feedbacks, e.g. growth resulting stress (Sampathkumar et al., 2014), auxin gradients (due to PIN; Shi et al., (2018)), soil thickness in roots, and bending through wind in the stem, there are bound to be feedback loops of cells and tissues to sense and react to those cues on a topological level to integrate this information into the plants development.

## Material and Methods

### Plant materials and growth condition

We grew *Arabidopsis thaliana* wild-type (WT; Wassilewskija ecotype) plants with the membrane reporter pUBQ10::acyl-YFP (previously described in (Willis et al., 2016)) and katanin1-2 mutant in Columbia-0 background with the membrane reporter Lti6b-GFP (Eng et al., 2021) in short day (8 h/16 h day/night), 20 °C/16 °C conditions for 3 weeks and then transferred to long day (16 h/8 h day/night), 20 °C/16 °C conditions till shoot apical meristem sampling. We cultured sampled shoot apical meristems (SAMs) in transparent imaging boxes containing apex culture media under long day, 22 °C conditions as previously described (Wang and Sampathkumar, 2020).

### Time-lapse data acquisition and pre-processing

We acquired confocal Z-stacks (3D images) at an excitation wavelength of 514 nm and 488 nm for imaging YFP and GFP respectively with a 40X/0.8 water immersion objective every 24 hours for 5 days (WT) or 3 days (*ktn*). Next, we used MorphoGraphX (MGX) (Barbier de Reuille et al., 2015) to obtain 2 ½ D surface mesh of the meristem L1 layer from the 3D images and from there we extract the cellular connectivity network (topology). In addition, we measured the shared cell wall of the neighboring cells (MGX function: Mesh/Export/Save Cell Neighborhood 2D), the surface area, and cell positions (MGX: Mesh/Heat Map/Heat Map Classic). The cellular connectivity network is composed of nodes, representing the centroids of the extracted cells. Edges connect two nodes if the corresponding cells are adjacent to each other. We lineage-tracked all cells between 1-day time steps manually in MGX (Figure 1A). We refer to dividing cells, at time t (days), as parent cells and their descendants, at time t + 1 (days), as daughter cells. To select the cells for the downstream analysis, we first manually determined the cells closest to the center of the SAM surface, given by the highest curvature. To this end, we compared the positions of cells to the average position over all cells.

### Prediction of dividing cells

To predict cell division events of central and non-peripheral cells, we selected all cells in a radius of 30 μm around the center (Figure 1B). In such a way, we only analyzed central and exclude peripheral cells. We considered a cell as peripheral with respect to a connectivity network in case the graph induced by the adjacent nodes does not form a cycle. We then labelled each of the selected cells as dividing (label: 1) or non-dividing (label: -1) cell within 24 hours (one day). In addition, we determined six sets of features (see below; for unreduced sets see Supplementary Figure 1) for each cell.

Five of six feature sets are based on the entire tissue (i.e. including peripheral cells as well as cells outside of the central region from the cellular connectivity network) and consist of topological features for all central cells; the sixth set includes only the surface area of the central cell. While calculating the topological properties, we considered different scenarios for weighting the edges. In the case of the unweighted topology, we weighted all edges equally (edge weight = 1). For the area-induced topology, we used the inverse of the mean surface areas of the two adjacent cells as edge weight. For the wall-induced topology, we defined the edge weights as the inverse of the shared cell wall area between two cells. For the distance-induced topology, we determined the inverse of the distance between the centroid positions of two adjacent cells as the edge weight (Figure 1C). We calculated ten topological properties for each central cell and network scenario (see Supplementary Table 1). Furthermore, we considered topological properties based on the induced subgraph of the first neighborhood (see Supplementary Table 1). We estimated all properties in python 3.8.1 using the networkx 2.4 package.

To train the classifiers for prediction of division events between two successive time points, we split the WT data from the five plants into two data sets, a training-validation and a testing set with four and one plant, respectively, while keeping three *ktn* plants as a separate test set. As there are fewer dividing cells, their class is the minority class. As a result, we down-sampled the majority class of non-dividing cells to balance the two classes. We applied a support vector machine (SVM) with a Gaussian kernel to predict the occurrence of cell division events within 1 day. To this end, we used the six different feature sets, namely: the unweighted topological features (unweighted topology), all topological features combined (topo), the surface area (surface area), topological features and area (topo + area), as well as two reduced feature sets including only topological features with Pearson correlation coefficients with surface area smaller than 0.5 or 0.7 (denoted by r < 0.5 and r < 0.7) (Supplementary Figure 1A). We trained each classifier with the topological properties as features of the training-validation set using five-fold cross-validation.

To this end, we z-normalised ((X-mean)/std) the topological properties with the corresponding mean and standard deviation (std) for the train-validation and the WT test data sets, respectively. The *ktn* data is z-normalised using its mean and standard deviations. We estimated the hyperparameters on the training set using another five-fold cross-validation using grid search (sklearn 0.22.1, GridSearchCV) regularly spacing 50 hyper-parameters for each power of 10. We further tested the classifiers by retraining the SVMs on all training-validation data with newly selected parameters and applied them on the unseen test data. We quantified the performance of the classifiers by calculating five measures of performance, including: the accuracy, F1-score, true positive rate, false positive rate, and area under the curve (AUC) of the receiver-operator characteristic (ROC). For comparative analysis between two performance measures p1 and p2, we used the relative difference (100 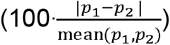

To further inspect the training of the classifiers, we generated the learning curves by retraining each classifier on a different number of training data (keeping the hyper-parameters from above). We further determined the feature sets information content by shuffling the labels 1000 times, retraining the classifier using the default RBF SVM parameters (sklearn 0.22.1, svm.SVC) on each set of shuffled labels, and calculating the performance of the resulting classifiers.

## Recreating of local topology after cell division

For the prediction of the changes in local topology of dividing cells, we selected all non-peripheral neighbor-parent-pairs of dividing cells. Next, we categorized the adjacency of these neighbors with respect to the newly divided (daughter) cells. To this end, we inspected if the neighbor of a neighbor-parent pair is adjacent to only one or both of the daughter cells.

To automate the procedure, we distinguished the divided daughter cells into the daughter closer to the center of the SAM which we termed cell “A” and the second daughter cell we named cell “B”. We labelled each neighbor cell in a neighbor-parent-pair with class 0, 1, or 2 according to whether it is connected only to cell “A”, cell “B”, or both. We then predicted the local topology excluding and including dividing neighbors using six feature sets, similar to the analysis above.

To distinguish neighbor-parent-pairs which are adjacent to two dividing cells, we considered the difference of topological features between neighbor and dividing parental cells in addition to the parental topological properties as features (Supplementary Figure 1B). As a result, we obtained the following feature sets: unweighted topology, topological features from all weightings (topo), biological features (bio, consisting of surface area and perimeter from neighbor and parent, as well as the shared cell wall and distance between the two), the combination of all topological and biological features (topo + bio), and two reduced feature sets (r < 0.5 and r < 0.7) including only topological features with Pearson correlation coefficients of smaller than 0.5 or 0.7 with all biological features. We performed training, validation, and testing as well as inspected the learning curves and estimated the information content of the used features as specified in the analysis above for wild type data. Additionally, we tested the classifiers on the *ktn* data.

### Application of the classifiers for division event and local topology

To combine the predictions of division events and local topology changes, we used the previously developed classifiers and applied them to predict how the topology of the test plants would change. To this end, we selected the classifiers including both topological and biological features (based on validation performance) and applied them one after another on to the test tissues to generate the topology of the next time points. Here, the predictions were only made for one time step (24 hours), since longer periods required us to estimate changes in the biological features as cells predicted to divide would not necessarily divide in the observed tissue one step later.

To arrive at the predicted cellular connectivity network, we determined the cells predicted to divide and divided cells’ future adjacency with their neighbors. Next, we repeated the following four steps for all cells predicted to divide at time t, starting with a random cell: (1) We removed the dividing cell along with the edges connecting the neighbors that is dividing. (2) We added the daughter cells representing the cell closer (cell A) and farther (cell B) away from the SAM center. (3) We connected the daughter cells with their neighbors based on the prediction from the local topology classifier (Figure 4A).

To evaluate the performance of the combined application of division and topology prediction, we calculated all unweighted topology features for the cells which are neither dividing in the predicted nor in the observed topology. Next, we plotted the non-dividing cells observed against the predicted features, determined best linear fit, and the Pearson correlation coefficient of all unweighted topological properties. In addition, we divided all predicted non-dividing cells, randomly assigned labels to the neighbors of dividing cells how they will be connected to the divided cells based on the training-validation set representation. Then, we repeated the correlation analysis from above 1000 times (differently reconnecting topologies), and compared the correlations between predicted and random topology propagation.

To also evaluate the local recreation of the topology around dividing cells, we compare the first neighborhoods of cells dividing in the predicted and observed tissue of the test plant by calculating the percentage of correctly labelled neighbors. The distributions of predicted accuracies are compared with an estimated random labelling of the neighbors using Kolmogorov-Smirnov-Test (scipy 1.4.1, ks_2samp).

## Code availability

The entire code to reproduce the findings is available at https://github.com/matz2532/SAM_division_prediction

## Acknowledgments

Z.N., T.M., A.S., Y.W., and R.K. acknowledge the support by the project SHAPENET, 031L0177A (to Z.N.) and 031L0177B (to A.S.), of the German Federal Ministry of Education and Research.

## Author Contributions

Z.N. and A.S. designed the research. T.W.M. implemented the computational approaches and performed the computational experiments. Y.W. and T.W.M. performed segmentation. Y.W. and R.K. performed the image generation. T.W.M. and Z.N. prepared the manuscript. All co-authors contributed to the final version of the manuscript.

## Supplementary Information

**Supplementary Figures 1.**
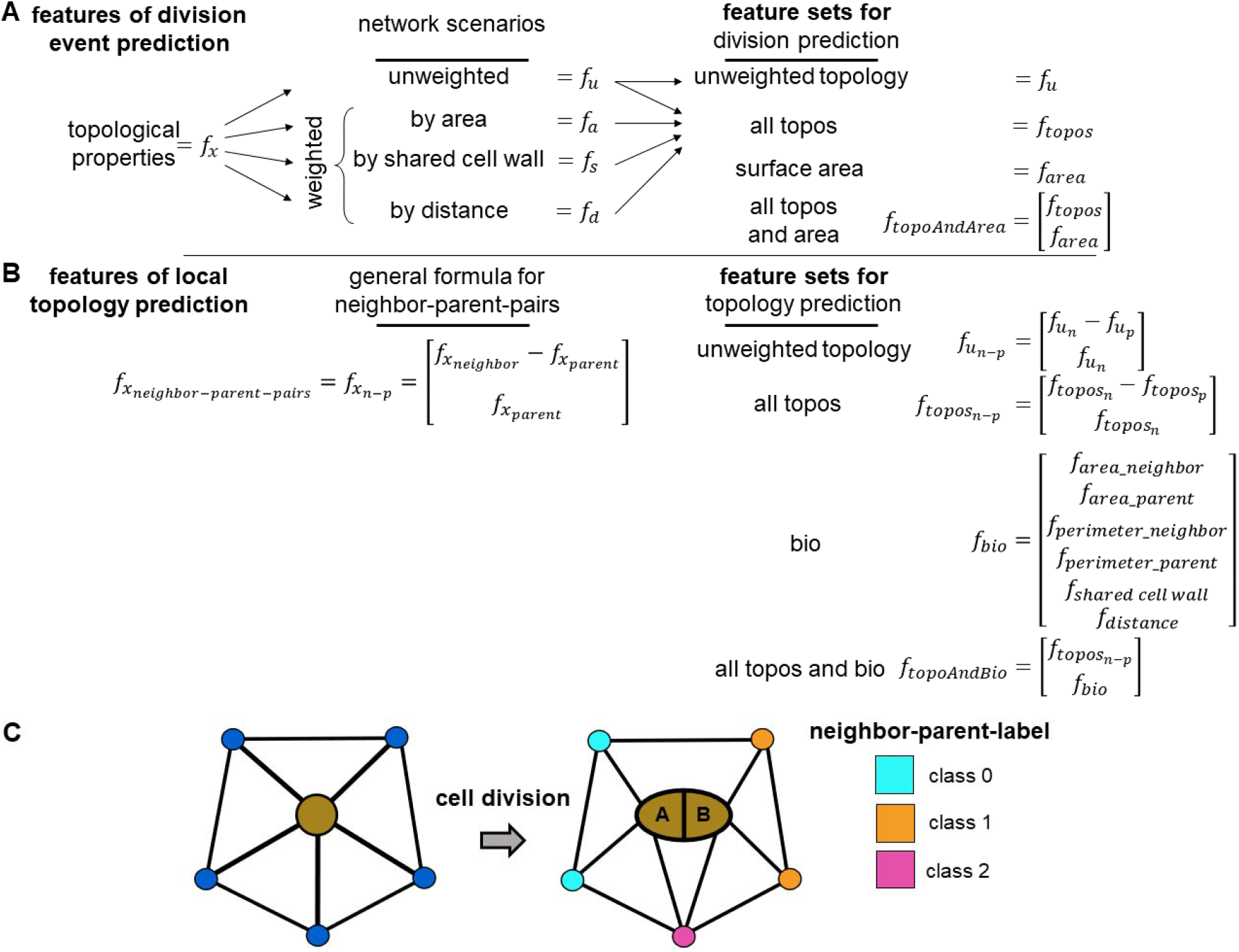
Overview about feature sets for division event and local topology prediction as well as an example for local topology class assignment. (A) For division event prediction, we consider 16 topological features (Supplementary Table 1) calculated from four network scenarios (unweighted, weighted by area, shared cell wall, and distance), creating four different feature sets: unweighted topology, all topologies (all topos), surface area, as well as topological features and surface area combined (all topos and area) for all central cells.(B) For local topology prediction, we calculate features for each neighbor-parent-pair of dividing parent cells using the difference in topological features of the neighbor and parent features as well as the features of the neighbor. Using this general formula, we generated four feature sets: unweighted topology, all topos, biological features (bio, including surface area, perimeter, shared cell wall, and distance), as well as topological and biological features combined (all topos and bio). (C) Parent cell (brown circle) divides into two daughter cells (A, B: representing the cell closer and farther away from the SAM center) changing the local topology in the process. The colors of the neighbors after division of the central cell represents the adjacency of the neighbor with the daughter cells: class 0 (cyan) neighbor is adjacent to cell A, class 1 (orange) pair neighbor is adjacent cell B, and class 2 (magenta) neighbors are adjacent to both cells. The classes are then used to be predicted from the earlier time point.

**Supplementary Figures 2.**
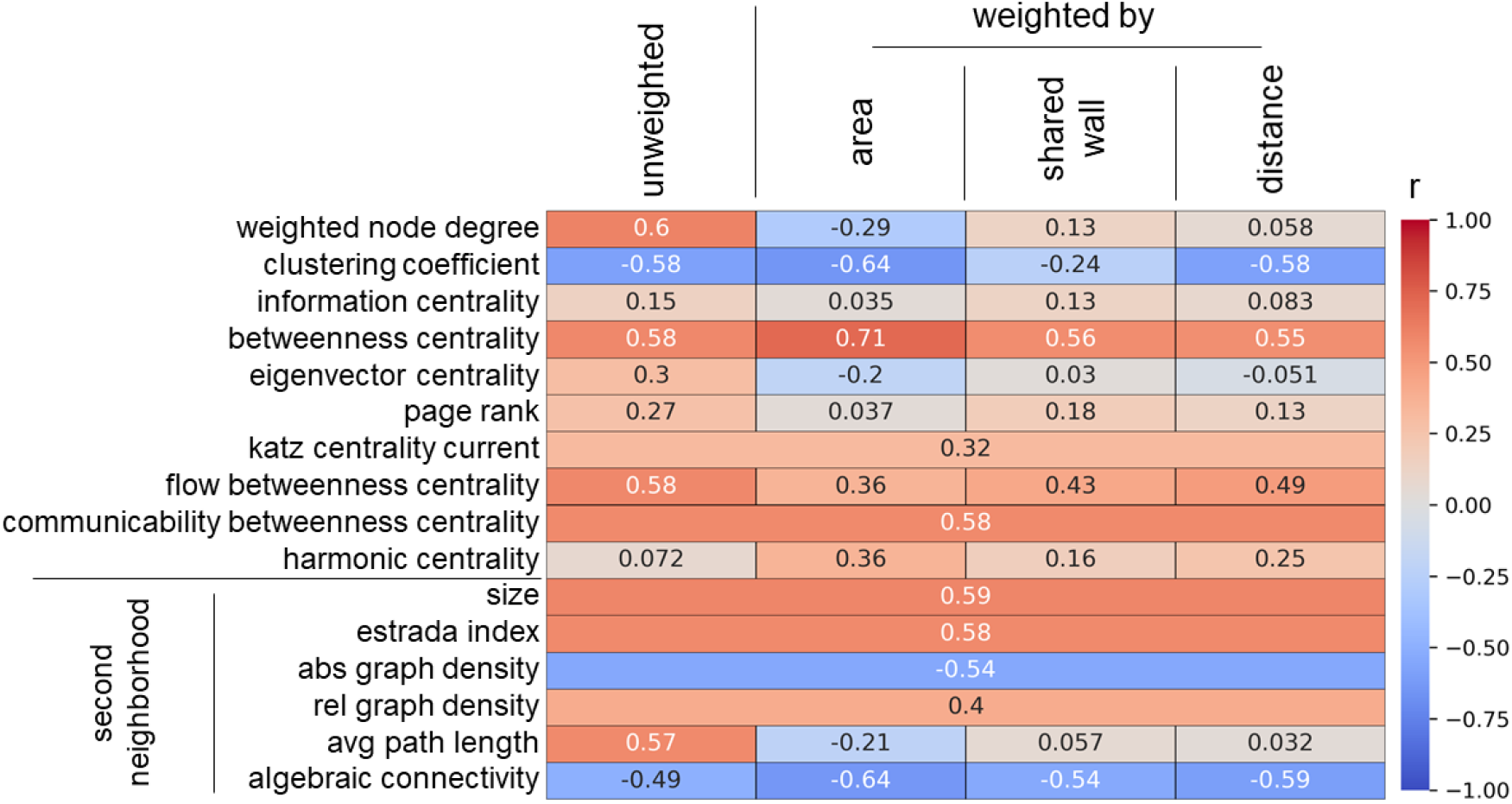
Heat map of Pearson correlation coefficients between topological features and surface area. We consider the 16 topological features calculated from the four network scenarios (see Figure 1C): unweighted edges and edges weighted by area, shared wall, and distance. Majority of topological features exhibit small Pearson correlation coefficients (r, legend range from -1 (blue) to 1 (red)). N_WT_ = 5 plants, 4 time steps, n_WT_ = 1225 cells.

**Supplementary Figure 3.**
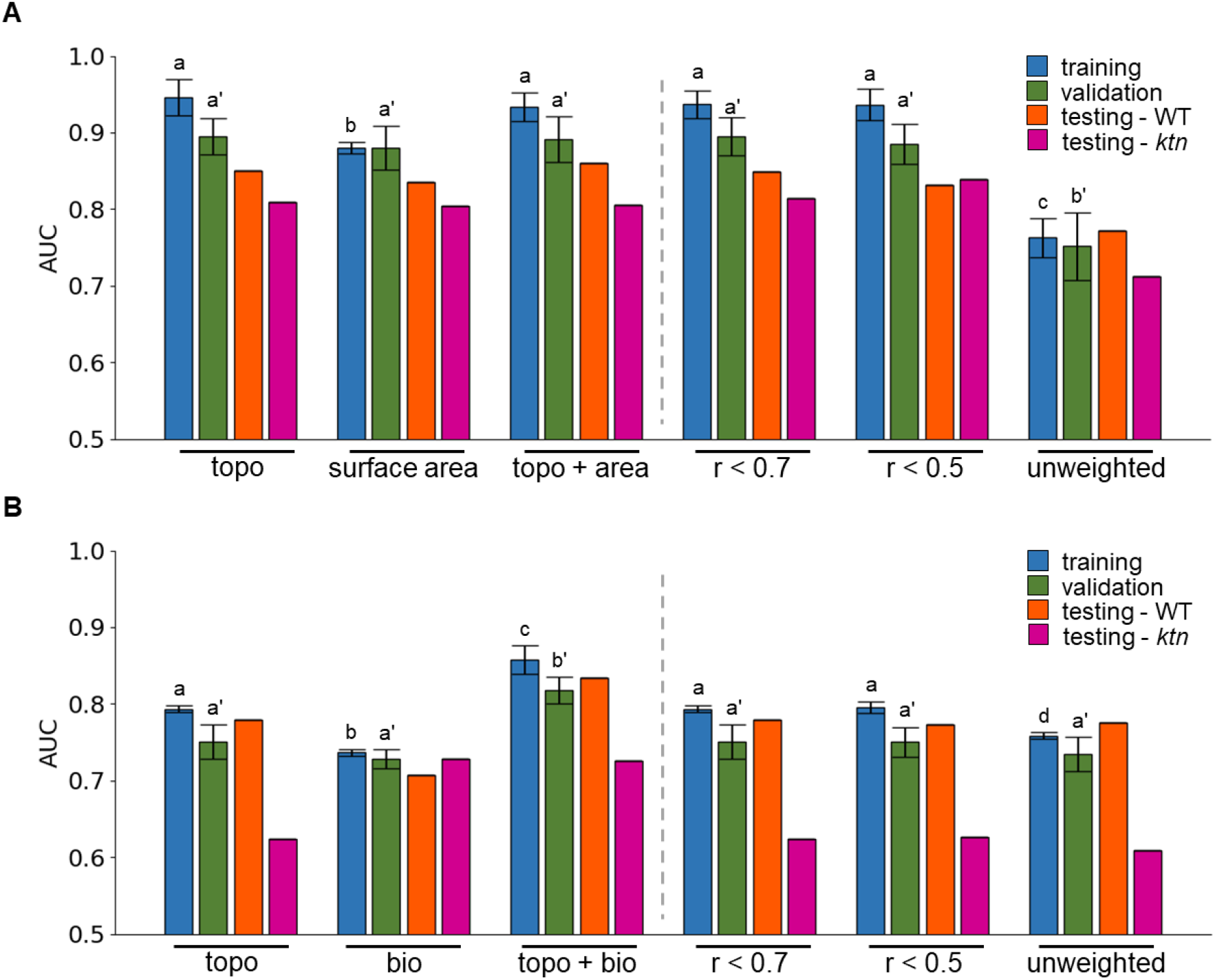
Comparative analysis of predictions based on reduced sets of features. Area under the curve (AUC) of the ROC of the support vector machine (SVM) classifier on the training (blue), validation (green), and testing of wild type (orange) and ktn mutant (purple) set of (A) division event and (B) local topology prediction. SVMs are trained on the combined topological features (topo), (A) surface area or (B) (bio, including surface area, perimeter, shared cell wall, and distance), topological features with (A) surface area (topo + area) or (B) bio (topo + bio), reduced set of topological features that show an absolute Pearson correlation coefficient with (A) surface area or (B) bio smaller than 0.7 or 0.5 (r < 0.7 and r < 0.5), as well as only the topological features derived from the unweighted network scenario (unweighted). The performance on the training and validation set is determined from five-fold cross-validation with mean and the standard deviation shown as error bars. Different letters indicate significance between groups using one-way ANOVA with Tukey’s pairwise comparison (p-value < 0.05). Statistical testing for differences of classifier performance for the training and validation sets was conducted separately (letter without and with apostrophe, respectively). N_WT_ = 5 plants, 4 time steps (4 plants for training-validation and 1 plant for testing); N_ktn_ = 3 plants, 3 time steps; (A) n_WT_ = 502 and 156, train-validation and test cells, n_ktn_ = 334 (balanced data) and (B) n_WT_ = 1317 and 321, train-validation and test cells, n_ktn_ = 888 (balanced data).

**Supplementary Figure 4.**
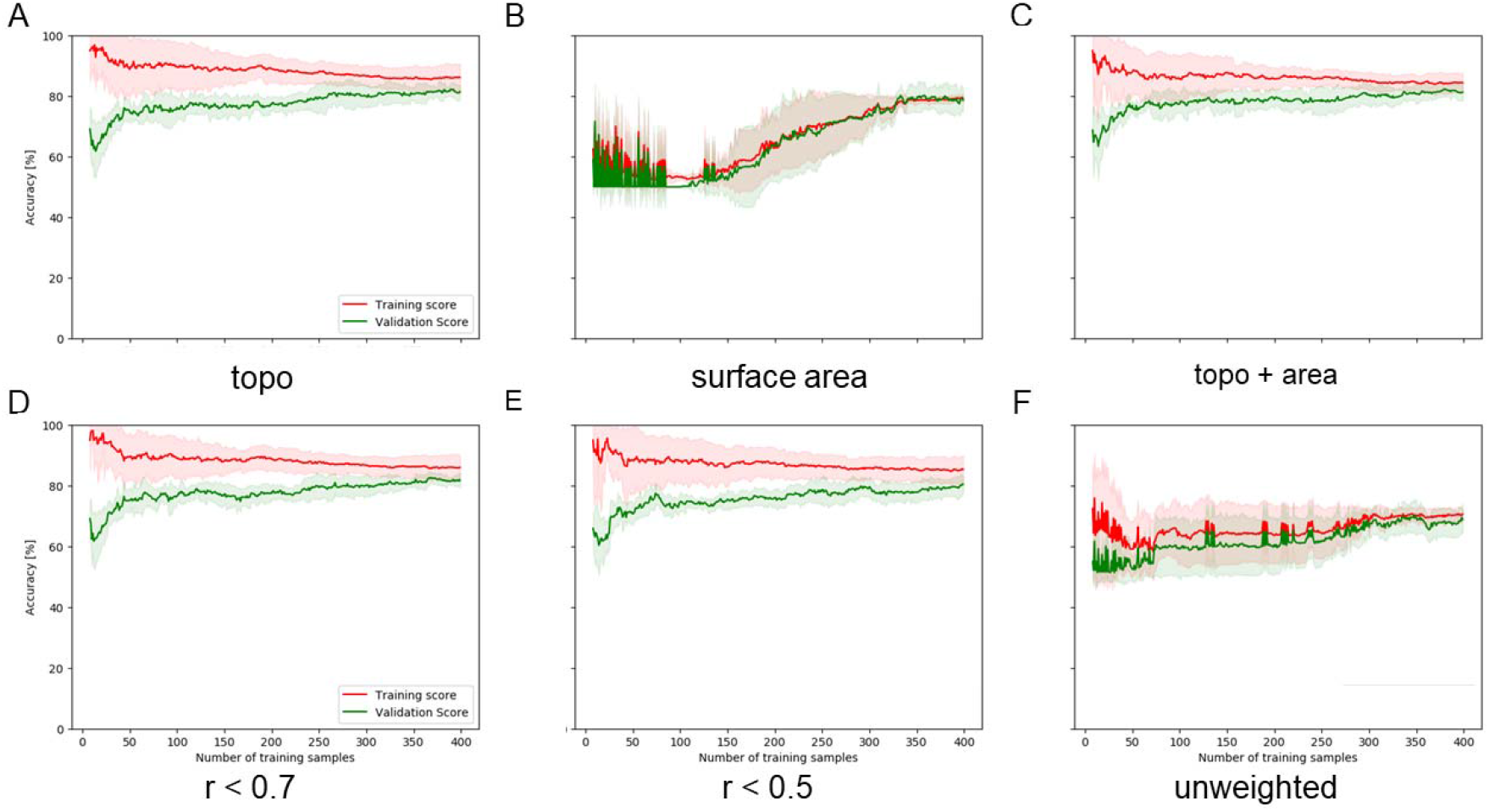
Learning curves for the classifiers that predict division events. Learning curves of SVMs predicting cell division events, based on four different feature sets, showing the accuracy in the validation (green) and training (red) set (line: mean, area: ± 1 standard deviation). Feature sets: (A) combined topological features (topo; including features calculated from the four network scenarios, see Figure 1C), (B) surface area as a single feature, topo with surface area (topo + area), (D, E) topological features which have an absolute Pearson correlation coefficient (r) with surface area smaller than 0.7 and 0.5, respectively, and (F) unweighted topological features (unweighted topology). N_WT_ = 4 plants, 4 time steps; n_WT_ = 502 train-validation cells (balanced data), one-way ANOVA.

**Supplementary Figure 5.**
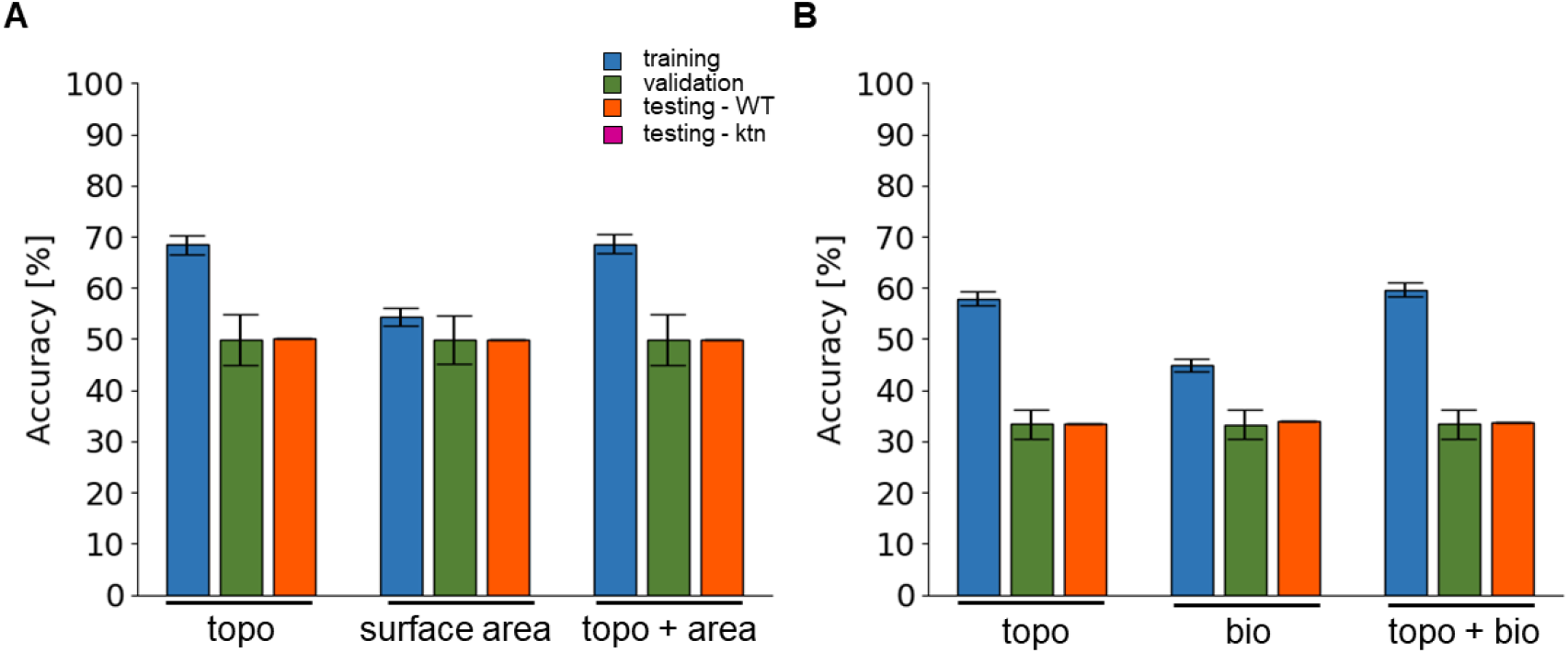
Performance of classifiers trained on randomized labels. Support vector machines (SVMs) trained on surface area/biological features and unweighted topological features perform similarly with respect to the prediction of a division event and local topology of shoot apical meristems (SAM). Accuracy of the SVM classifier on the training (blue), validation (green), and testing (orange) set for a division event (A) and local topology prediction (B) based on unweighted topological features (unweighted topology), surface area, or biological features (bio). Shown are the mean and standard deviation on the training and validation sets from five-fold cross-validation. The performance on the training and validation set is determined from five-fold cross-validation with mean and the standard deviation shown as error bars. Different letters indicate significance between groups using one-way ANOVA with Tukey’s pairwise comparison: p-value < 0.05. Statistical testing for differences of classifier performance for the training and validation sets was conducted separately (letter without and with apostrophe, respectively). N_WT_ = 5 plants, 4 time steps (4 plants for training-validation and 1 plant for testing); (A) n_WT_ = 502 and 156, and (B) n_WT_ = 1317 and 321 train-validation and test cells respectively (balanced data) shuffling labels 1000 times.

**Supplementary Figure 6.**
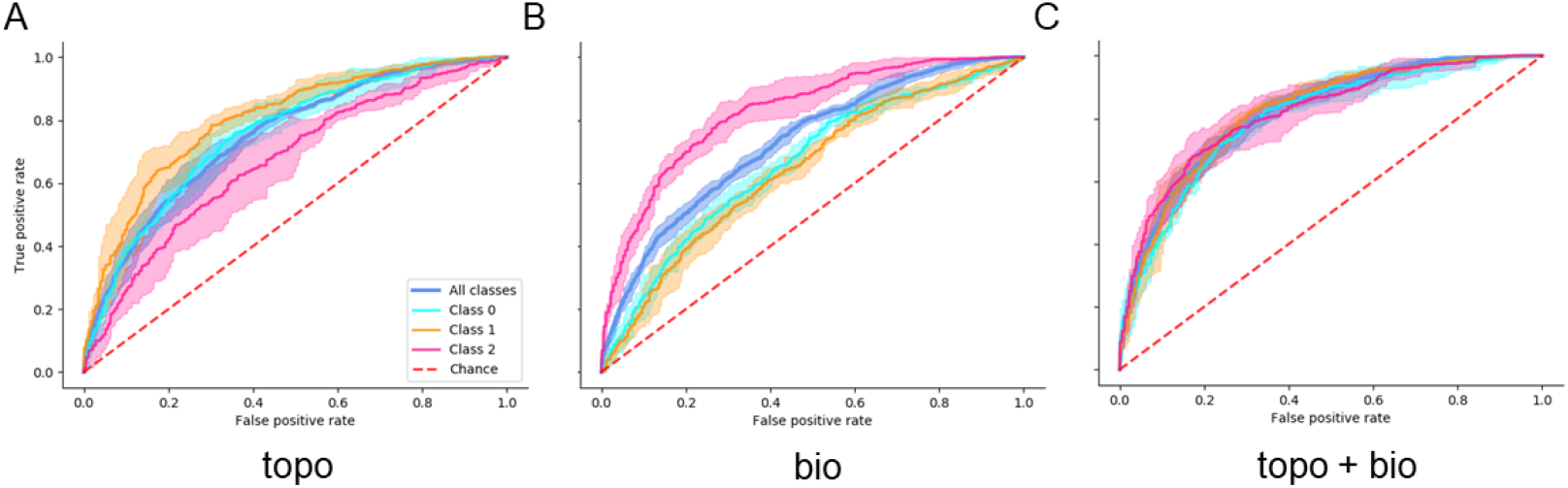
Difference between classifiers trained on topological and biological features to predict local topology. Classifiers based on topological and biological features predict the three different classes of cells that are neighbors of cells that have divided in comparison to the previous time point. Receiver operating characteristic-curve predicting cell division events on five-fold cross-validation of (A) combined topological features (topo), including features calculated from the four network scenarios (see Figure 1C), (B) biological features (bio), including surface area, perimeter, shared cell wall, and distance, and (C) topological and biological features combined (topo + bio). The mean performance is shown as a straight line, together with the area of ± 1 standard deviation obtained from the five-fold cross validation of the average ROC-curve combining all classes (blue), of class 0 (cyan), class 1 (orange), and class 2 (magenta). The performance expected by change is marked with a red dashed line. N_WT_ = 4 plants, 4 time steps; n_WT_ = 1317 train-validation cells (balanced data).

**Supplementary Figure 7.**
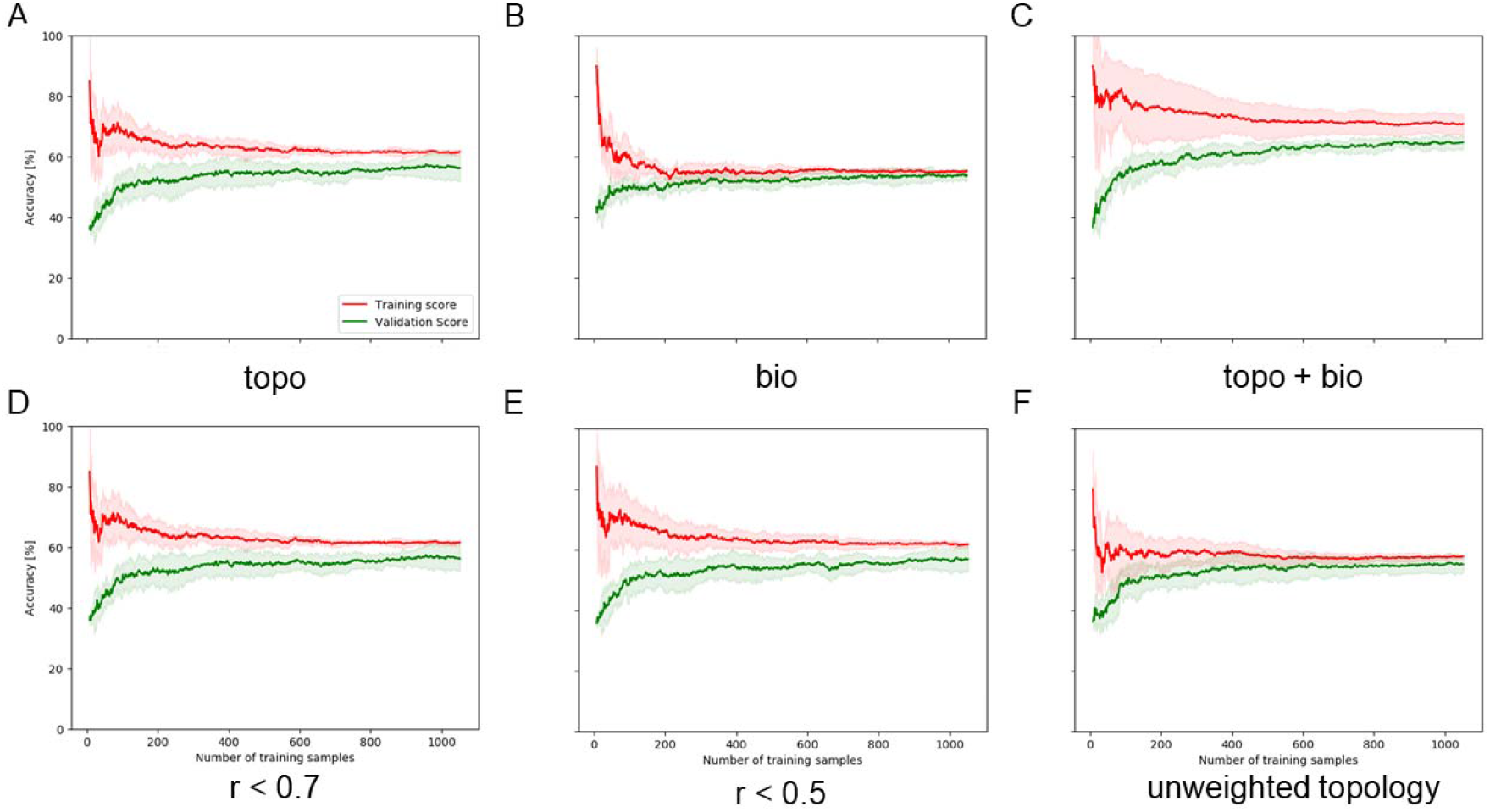
Learning curves for the classifiers predicting local topological changes. Learning curves of support vector machines (SVMs) that predict local topology based on six different feature sets: (A) combined topological features (topo, including features calculated from the four network scenarios, see Figure 1C), (B) biological features (bio), including surface area, perimeter, shared cell wall, and distance, (C) topological and biological features combined (topo+bio), (D, E) topological features which have an absolute value of Pearson correlation coefficients with all biological features (cor) smaller than 0.7 or 0.5, and (F) only unweighted features showing validation (green) and training (red) accuracy (line: mean, area: ± 1 standard deviation). N_WT_ = 4 plants, 4 time steps; n_WT_ = 1317 train-validation cells (balanced data).

**Supplementary Figure 8.**
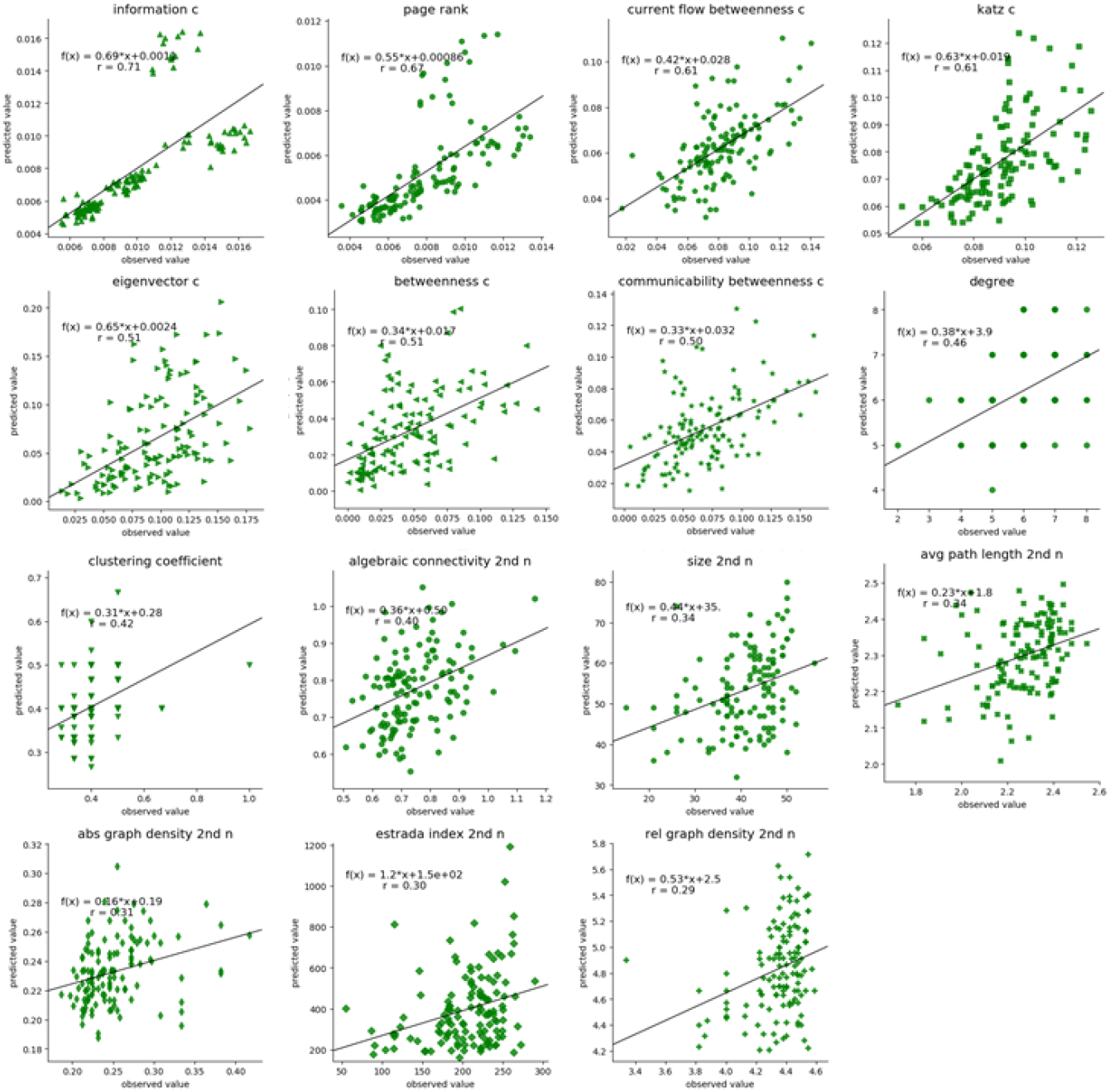
Observed and predicted features from non-dividing cells plotted against each other. The observed feature values are plotted against the predicted features values for each cell non-dividing in the observed and predicted tissue. A line fitting the linear regression line is plotted to the data including its function f(x) and Pearson correlation coefficient r for the unweighted topological features: information c (centrality), page rank, current flow betweenness c, katz c, communicability betweenness c, load c, betweenness c, eigenvector c, clustering coefficient, degree, avg. path length on 2 neighborhood (2nd n), abs graph density 2nd n, size 2nd n, algebraic connectivity 2nd n, rel. graph density 2nd n, estrada index 2nd n. The predicted tissue is estimated using the division and topology prediction classifiers trained on topological and biological features of the training-validation data (see Figure 4E). N_WT_ = 1 plant (test), 4 time steps; n_WT_ = 126.

**Supplementary Figure 9.**
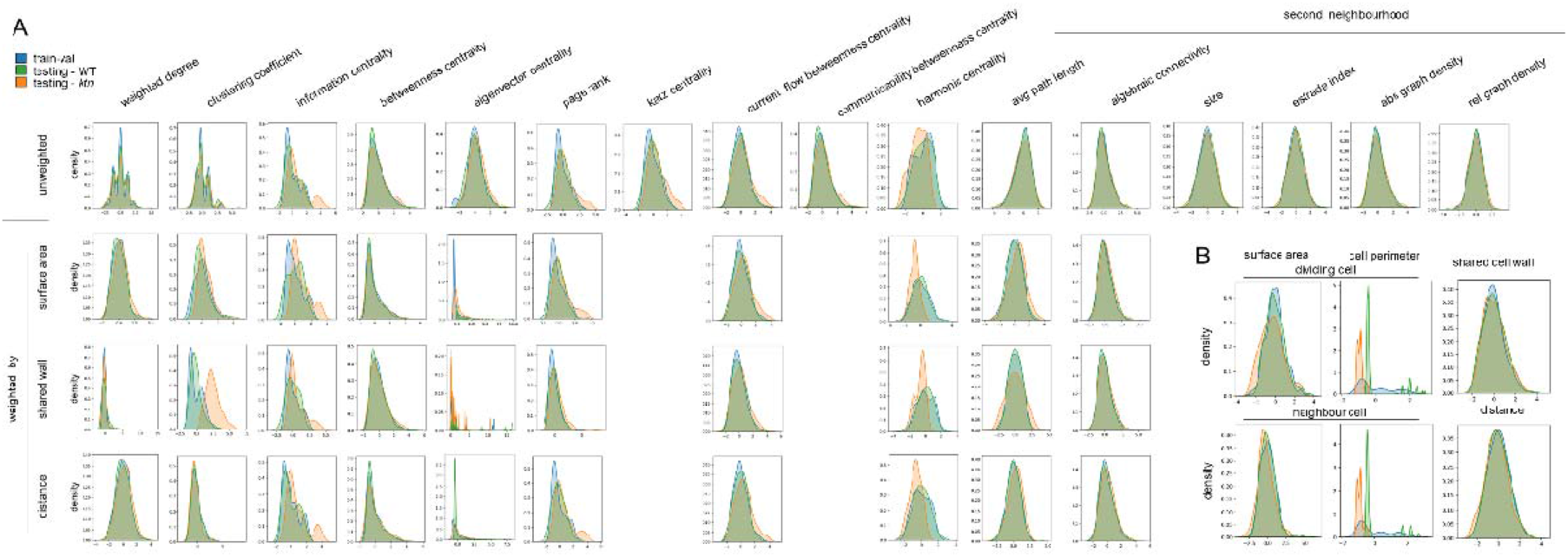
Density distributions of topological and biological features of WT and ktn data. The density distributions of the normalized (A) topological and (B) biological features are plotted to differentiate between pooled samples used during training and validation from wild-type (WT; blue), and testing from WT (green), and ktn (orange). The topological features are grouped into unweighted and weighted by area, shared cell wall, and distance (see Figure 1C), while topological features independent of the network scenario are only displayed in the unweighted row. The columns are further grouped into features calculated on the second neighborhood and features from the dividing cell or its neighbor. (A) n_WT_ = 502, 156, train-validation and test cells respectively; n_ktn_ = 334 (balanced data) (B) n_WT_ = 1317 and 312, train-validation and test cells respectively; n_ktn_ = 888 (balanced data).

**Supplementary Table 1.** Definition of network centralities applied on differently weighted network scenarios to be used as feature sets for cell division and local topology prediction. Each network centrality was applied on the unweighted, weighted by area, shared cell wall, and distance network scenario. The network centralities were concatenated together for each cell and used to train and predict different classifiers.

| Network centrality | Definition | Reference |
| --- | --- | --- |
| weighted node degree | $C_D(v) = \frac{\deg(v)}{\vert V\vert - 1}$ | Freeman, 1979 |
| clustering coefficient | $C_u = \frac{1}{\deg(u)(\deg(u) - 1)} \sum_{vw} \hat{w}_{uv} \hat{w}_{uw} \hat{w}_{vw}$ | Saramäki, 2007 |
| information centrality | $C_I(v) = \frac{\vert V\vert - 1}{\sum_{u \in V} p_{uv}(v) p_{uv}(u)}$ | Sabidussi, 1966 |
| betweenness centrality | $C_B(v) = \frac{2}{(\vert V\vert - 1)(\vert V\vert - 2)} \sum_{s, t \in V} \frac{q_v(s, t)}{q(s, t)}$ | Freeman, 1977 |
| eigenvector centrality | $C_E(v) = \frac{1}{\lambda} \sum_{u \in V} a_{vu} C_E(u)$ | Bonacich, 1987 |
| PageRank | $R'(u) = c \sum_{v \in B_u} \frac{R'(v)}{N_v} + cE(u)$ | Page, 1999 |
| Katz centrality | $x_i = \alpha \sum_j A_{ij} + \beta$ | Katz, 1953 |
| current flow betweenness centrality | $c_{CB}(v) = \frac{1}{(n - 1)(n - 2)} \sum_{s, t \in V} \tau_{st}(v)$ | Newman, 2005 |
| communicability betweenness centrality | $c_{CFS}(v) = \frac{2}{(\vert V\vert - 1)(\vert V\vert - 2)} \sum_{s, t \in V} \tau_{st}(v)$ | Newman, 2005 |
| harmonic centrality | $C_H(v) = \frac{\sum_{u \in V} \frac{1}{d(u, v)}}{\vert V\vert - 1}$ | Marchiori & Latora, 2000 |
| size | number of edges in a graph |  |
| Estrada index | $EEG(G) = \sum_{j=1}^n e^{\lambda_j}$ | Estrada, 200 |
| absolute graph density | $C = \frac{2m}{n(n - q)}$ | Wilks & Meara, 2002 |
| relative graph density | $C = \frac{2m}{n}$ | |
| average path length | $a = \sum_{s, t \in V} \frac{d(s, t)}{n(n - 1)}$ | Mao & Zhang, 2013 |
| algebraic connectivity | second smallest eigenvalue of Laplacian matrix | Fiedler, 1973 |

**Supplementary Table 2.** Features ordered by Pearson correlation coefficient with area. Each network centrality was applied on the unweighted, weighted by area, shared cell wall, and distance network scenario. The Pearson correlation coefficient with area was calculated and were ordered from highest to lowest.

| Feature name | Pearson correlation coefficient | p-value |
| --- | --- | --- |
| betweenness centrality weighted by area | 0.711959318 | 4.858135677288066e-211 |
| degree | 0.603854449 | 4.04599560747419e-136 |
| size on 2 neighborhood | 0.592735441 | 5.437779671764918e-130 |
| betweenness centrality | 0.583915039 | 2.6889326200117693e-125 |
| current flow betweenness centrality | 0.580737704 | 1.217610917275356e-123 |
| load centrality | 0.579975133 | 3.0215367111277033e-123 |
| estrada index on 2 neighborhood | 0.577117774 | 8.912704904812637e-122 |
| estrada index on 2 neighborhood weighted by area | 0.577117774 | 8.912704904812637e-122 |
| communicability betweenness centrality | 0.57662665 | 1.5891934730974062e-121 |
| communicability betweenness centrality weighted by area | 0.57662665 | 1.5891934731075239e-121 |
| avg path length on 2 neighborhood | 0.571341971 | 7.534891764175505e-119 |
| betweenness centrality weighted by shared wall | 0.562314483 | 2.17398436729873e-114 |
| betweenness centrality weighted by distance | 0.551932822 | 1.9924782594347459e-109 |
| current flow betweenness centrality weighted by distance | 0.49002158 | 3.548421338397669e-83 |
| current flow betweenness centrality weighted by shared wall | 0.425038752 | 7.271052967178761e-61 |
| rel graph density on 2 neighborhood | 0.402000467 | 4.72658491702921e-54 |
| current flow betweenness centrality weighted by area | 0.357475281 | 2.5263809251463147e-42 |
| harmonic centrality weighted by area | 0.356480643 | 4.4059338421761555e-42 |
| katz centrality weighted by area | 0.318100276 | 2.1117164600213396e-33 |
| katz centrality | 0.318100276 | 2.1117164600213396e-33 |
| eigenvector centrality | 0.296303239 | 5.28165100633828e-29 |
| page rank | 0.268477018 | 6.447099545102352e-24 |
| harmonic centrality weighted by distance | 0.247926272 | 1.5906836425501315e-20 |
| page rank weighted by shared wall | 0.181567285 | 1.4700578049314804e-11 |
| harmonic centrality weighted by shared wall | 0.164231644 | 1.0815009609397551e-09 |
| information centrality | 0.14740273 | 4.633731043029875e-08 |
| information centrality weighted by shared wall | 0.13173255 | 1.0692961604944774e-06 |
| weighted node degree weighted by shared wall | 0.128973752 | 1.7935045888103348e-06 |
| page rank weighted by distance | 0.127130861 | 2.5188037924397987e-06 |
| information centrality weighted by distance | 0.082746096 | 0.002241568 |
| harmonic centrality | 0.071745324 | 0.008078663 |
| weighted node degree weighted by distance | 0.057524217 | 0.033774282 |
| avg path length on 2 neighborhood<br>weighted by shared wall | 0.056763189 | 0.03620415 |
| page rank weighted by area | 0.037413208 | 0.167598817 |
| information centrality weighted by area | 0.035267233 | 0.19334169 |
| avg path length on 2 neighborhood<br>weighted by distance | 0.0323718 | 0.232514552 |
| eigenvector centrality<br>weighted by shared wall | 0.03031164 | 0.263614219 |
| eigenvector centrality weighted by distance | -0.051181347 | 0.05897689 |
| eigenvector centrality weighted by area | -0.203077703 | 3.8271343446663024e-14 |
| avg path length on 2 neighborhood<br>weighted by area | -0.214150561 | 1.361632834653018e-15 |
| clustering coefficient<br>weighted by shared wall | -0.236129215 | 1.0319617716515097e-18 |
| weighted node degree weighted by area | -0.289769264 | 9.320458571568519e-28 |
| algebraic connectivity on 2 neighborhood | -0.493010144 | 2.5317849899442773e-84 |
| algebraic connectivity on 2 neighborhood<br>weighted by shared wall | -0.535130612 | 9.326326866241923e-102 |
| abs graph density on 2 neighborhood | -0.54315277 | 2.3001423114310698e-105 |
| clustering coefficient | -0.580538544 | 1.544179245441756e-123 |
| clustering coefficient weighted by distance | -0.584348465 | 1.5932034339322458e-125 |
| algebraic connectivity on 2 neighborhood<br>weighted by distance | -0.594101302 | 9.896429835985306e-131 |
| algebraic connectivity on 2 neighborhood<br>weighted by area | -0.639772476 | 1.1317137430247356e-157 |
| clustering coefficient weighted by area | -0.642874421 | 1.1369324719240177e-159 |

**Supplementary Table 3.** Performance measures of division event prediction from SVMs trained on features from four differently weighted topologies and/or area. F1-score (F1), accuracy (Acc), true positive rate (TPR), false positive rate (FPR), and area under the ROC-curve (Auc) of training (train) and unseen data (val) on the splits of the five-fold cross-validations, as well as retraining on train+val data testing on test data (testing), their mean and standard deviation (std) as well as the performance of testing on a never seen plant training on the full training-validation data set. Feature sets: combined topological features (topo; including features of unweighted, weighted by area-, shared cell wall-, and distance topologies), area as a single feature, topo with area, topological features which have an absolute Pearson correlation coefficient with area smaller than 0.7 or 0.5 (cor < 0.7 and cor < 0.5, respectively) and unweighted topological features (unweighted topology).

topo
|  | train F1 | train Acc | train TPR | train FPR | train Auc | val F1 | val Acc | val TPR | val FPR | val Auc |
| --- | --- | --- | --- | --- | --- | --- | --- | --- | --- | --- |
| split 0 | 91.36 | 91.25 | 92.50 | 10.00 | 0.9697 | 82.69 | 82.35 | 84.31 | 19.61 | 0.9250 |
| split 1 | 85.31 | 84.58 | 89.55 | 20.40 | 0.9133 | 80.00 | 80.00 | 80.00 | 20.00 | 0.8708 |
| split 2 | 93.63 | 93.53 | 95.02 | 7.96 | 0.9702 | 78.85 | 78.00 | 82.00 | 26.00 | 0.8724 |
| split 3 | 86.46 | 85.82 | 90.55 | 18.91 | 0.9244 | 84.62 | 84.00 | 88.00 | 20.00 | 0.9220 |
| split 4 | 88.45 | 88.31 | 89.55 | 12.94 | 0.9511 | 84.62 | 84.00 | 88.00 | 20.00 | 0.8836 |
| mean | 89.04 | 88.70 | 91.44 | 14.04 | 0.9457 | 82.15 | 81.67 | 84.46 | 21.12 | 0.8948 |
| std | 3.08 | 3.33 | 2.09 | 4.87 | 0.0233 | 2.37 | 2.35 | 3.19 | 2.44 | 0.0239 |
| testing | 82.17 | 81.67 | 84.46 | 21.12 | 0.8922 | 76.62 | 76.92 | 75.64 | 21.79 | 0.8498 |
| ktn testing |  |  |  |  |  | 69.93 | 72.46 | 64.07 | 19.16 | 0.8087 |

area
|  | train F1 | train Acc | train TPR | train FPR | train Auc | val F1 | val Acc | val TPR | val FPR | val Auc |
| --- | --- | --- | --- | --- | --- | --- | --- | --- | --- | --- |
| split 0 | 76.60 | 78.00 | 72.00 | 16.00 | 0.8674 | 80.00 | 81.37 | 74.51 | 11.76 | 0.9285 |
| split 1 | 78.43 | 78.11 | 79.60 | 23.38 | 0.8773 | 82.83 | 83.00 | 82.00 | 16.00 | 0.8940 |
| split 2 | 80.60 | 80.85 | 79.60 | 17.91 | 0.8873 | 74.51 | 74.00 | 76.00 | 28.00 | 0.8536 |
| split 3 | 78.97 | 79.60 | 76.62 | 17.41 | 0.8836 | 78.35 | 79.00 | 76.00 | 18.00 | 0.8664 |
| split 4 | 80.40 | 80.60 | 79.60 | 18.41 | 0.8864 | 75.00 | 76.00 | 72.00 | 20.00 | 0.8556 |
| mean | 79.00 | 79.43 | 77.48 | 18.62 | 0.8804 | 78.14 | 78.67 | 76.10 | 18.75 | 0.8796 |
| std | 1.46 | 1.20 | 2.98 | 2.51 | 0.0074 | 3.12 | 3.32 | 3.29 | 5.37 | 0.0284 |
| testing | 79.36 | 79.48 | 78.88 | 19.92 | 0.8804 | 69.06 | 72.44 | 61.54 | 16.67 | 0.8358 |
| ktn testing |  |  |  |  |  | 67.34 | 70.96 | 59.88 | 17.96 | 0.8046 |

topo with area
|  | train F1 | train Acc | train TPR | train FPR | train Auc | val F1 | val Acc | val TPR | val FPR | val Auc |
| --- | --- | --- | --- | --- | --- | --- | --- | --- | --- | --- |
| split 0 | 89.43 | 89.25 | 91.00 | 12.50 | 0.9594 | 83.81 | 83.33 | 86.27 | 19.61 | 0.9319 |
| split 1 | 85.37 | 84.83 | 88.56 | 18.91 | 0.9172 | 82.00 | 82.00 | 82.00 | 18.00 | 0.8804 |
| split 2 | 86.34 | 86.07 | 88.06 | 15.92 | 0.9307 | 80.00 | 79.00 | 84.00 | 26.00 | 0.8424 |
| split 3 | 83.25 | 82.59 | 86.57 | 21.39 | 0.9093 | 84.31 | 84.00 | 86.00 | 18.00 | 0.9072 |
| split 4 | 88.89 | 88.81 | 89.55 | 11.94 | 0.9498 | 82.35 | 82.00 | 84.00 | 20.00 | 0.8924 |
| mean | 86.66 | 86.31 | 88.75 | 16.13 | 0.9333 | 82.50 | 82.07 | 84.45 | 20.32 | 0.8909 |
| std | 2.28 | 2.49 | 1.48 | 3.64 | 0.0189 | 1.52 | 1.72 | 1.56 | 2.95 | 0.0297 |
| testing | 80.94 | 80.68 | 82.07 | 20.72 | 0.8935 | 77.85 | 78.85 | 74.36 | 16.67 | 0.8601 |
| ktn testing |  |  |  |  |  | 71.01 | 73.35 | 65.27 | 18.56 | 0.8056 |

**cor < 0.7**
|  | train F1 | train Acc | train TPR | train FPR | train Auc | val F1 | val Acc | val TPR | val FPR | val Auc |
| --- | --- | --- | --- | --- | --- | --- | --- | --- | --- | --- |
| split 0 | 85.57 | 85.25 | 87.50 | 17.00 | 0.9343 | 84.91 | 84.31 | 88.24 | 19.61 | 0.9239 |
| split 1 | 85.58 | 84.83 | 90.05 | 20.40 | 0.9182 | 81.19 | 81.00 | 82.00 | 20.00 | 0.8696 |
| split 2 | 93.37 | 93.28 | 94.53 | 7.96 | 0.9706 | 80.00 | 79.00 | 84.00 | 26.00 | 0.8688 |
| split 3 | 86.26 | 85.57 | 90.55 | 19.40 | 0.9253 | 85.44 | 85.00 | 88.00 | 18.00 | 0.9236 |
| split 4 | 86.47 | 86.07 | 89.05 | 16.92 | 0.9358 | 84.91 | 84.00 | 90.00 | 22.00 | 0.8884 |
| mean | 87.45 | 87.00 | 90.34 | 16.34 | 0.9369 | 83.29 | 82.66 | 86.45 | 21.12 | 0.8949 |
| std | 2.98 | 3.17 | 2.34 | 4.40 | 0.0181 | 2.24 | 2.29 | 2.97 | 2.75 | 0.0246 |
| testing | 81.78 | 81.27 | 84.06 | 21.51 | 0.8960 | 77.22 | 76.92 | 78.21 | 24.36 | 0.8488 |
| ktn testing |  |  |  |  |  | 69.13 | 72.46 | 61.68 | 16.77 | 0.8145 |

**cor < 0.5**
|  | train F1 | train Acc | train TPR | train FPR | train Auc | val F1 | val Acc | val TPR | val FPR | val Auc |
| --- | --- | --- | --- | --- | --- | --- | --- | --- | --- | --- |
| split 0 | 68.50 | 68.50 | 68.50 | 31.50 | 0.7510 | 72.00 | 72.55 | 70.59 | 25.49 | 0.7559 |
| split 1 | 75.41 | 73.88 | 80.10 | 32.34 | 0.8109 | 62.96 | 60.00 | 68.00 | 48.00 | 0.6796 |
| split 2 | 69.52 | 69.90 | 68.66 | 28.86 | 0.7602 | 69.31 | 69.00 | 70.00 | 32.00 | 0.7288 |
| split 3 | 69.44 | 68.91 | 70.65 | 32.84 | 0.7396 | 76.19 | 75.00 | 80.00 | 30.00 | 0.8012 |
| split 4 | 70.26 | 71.14 | 68.16 | 25.87 | 0.7509 | 65.93 | 69.00 | 60.00 | 22.00 | 0.7920 |
| mean | 70.63 | 70.47 | 71.21 | 30.28 | 0.7625 | 69.28 | 69.11 | 69.72 | 31.50 | 0.7515 |
| std | 2.46 | 1.94 | 4.53 | 2.60 | 0.0251 | 4.61 | 5.09 | 6.39 | 8.96 | 0.0443 |
| testing | 68.94 | 69.12 | 68.53 | 30.28 | 0.7512 | 70.66 | 68.59 | 75.64 | 38.46 | 0.7720 |
| ktn testing |  |  |  |  |  | 64.71 | 64.07 | 65.87 | 37.72 | 0.7115 |

**Supplementary Table 4.**
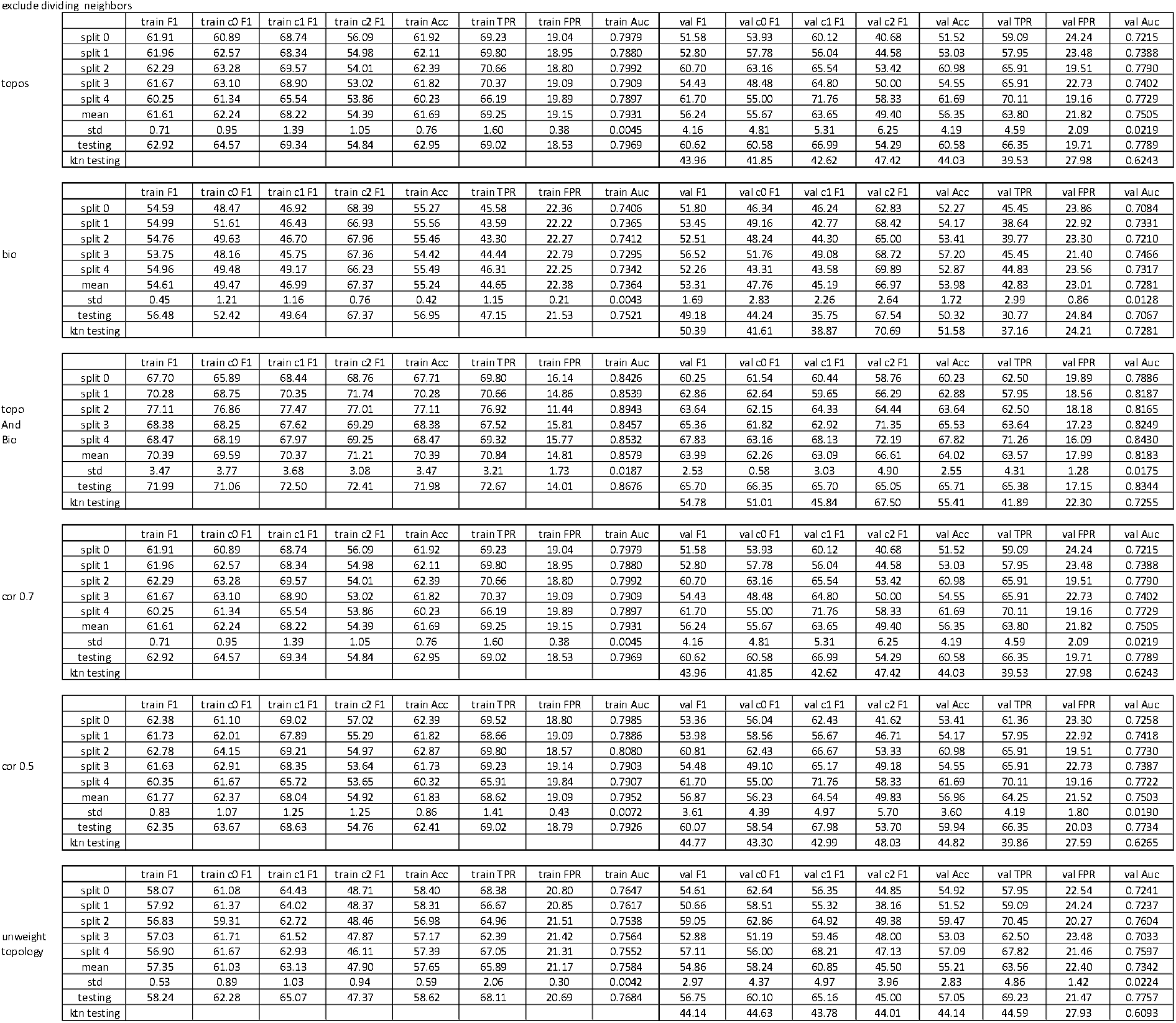
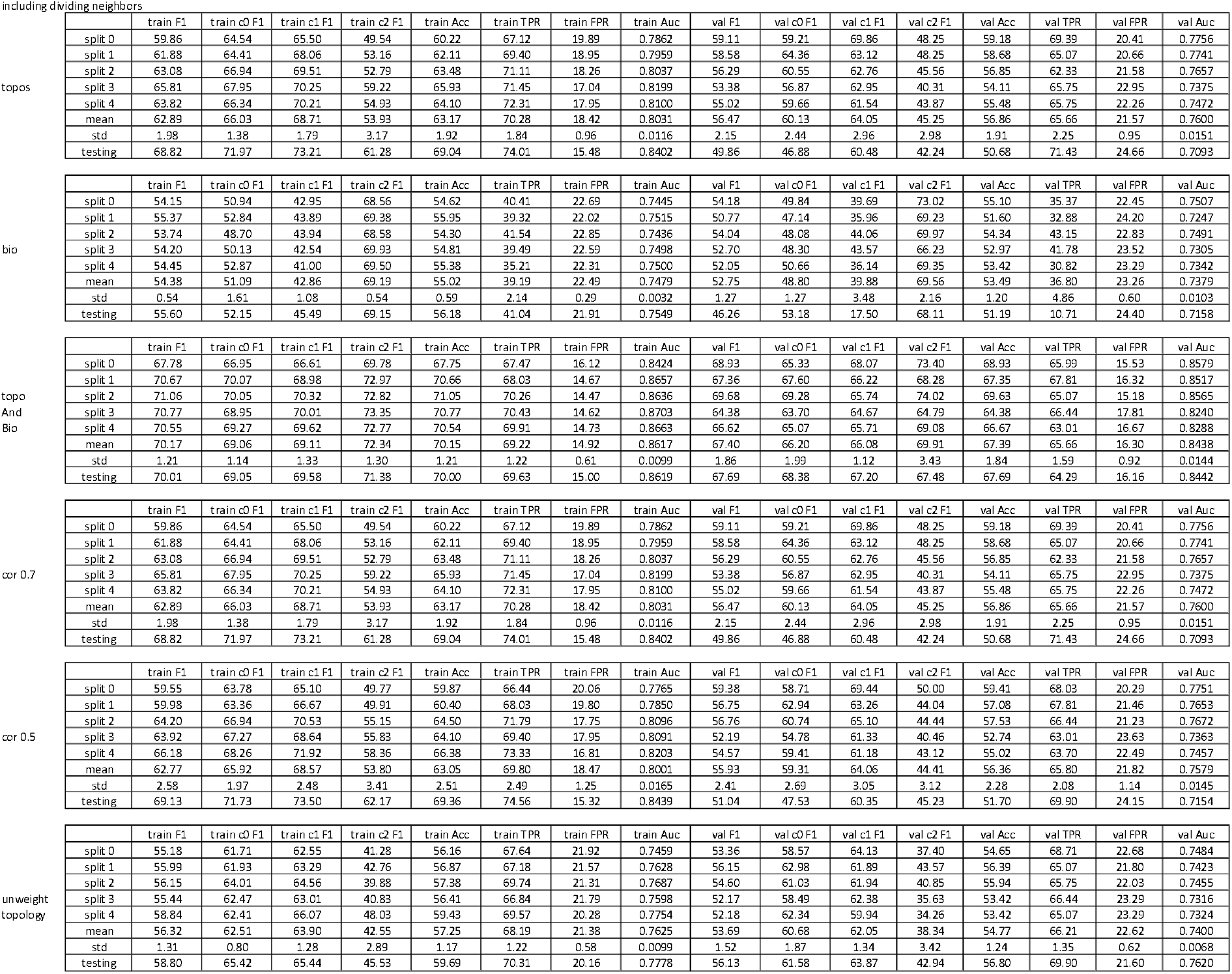
Performance measures of local topology prediction from SVMs trained from four different topologies and/or biological features. F1-score (F1), accuracy (Acc), true positive rate (TPR), false positive rate (FPR), and area under the ROC-curve (Auc) of training (train) and unseen data (val) on the splits of the five-fold cross-validations, as well as retraining on train+val data testing on test data (testing) (A) excluding or (B) including dividing neighbours, their mean and standard deviation (dev) as well as the performance of testing on a never seen plant training on the full training-validation data set. Feature sets: combined topological features (topo; including features of unweighted, weighted by area-, shared cell wall-, and distance topologies), biological features (bio; including area, perimeter, shared cell wall, and distance), topo with bio, topological features which have an absolute value of the Pearson correlation coefficients with all biological features (cor) smaller than 0.7 and 0.5, respectively, and unweighted topological features (unweighted topology).

